# Polycomb Marked Promoter Methylation Gains in Aging and Cancer are Independent of Replication

**DOI:** 10.64898/2026.10.30.763235

**Authors:** Sara-Jayne Thursby, Nibedita Patel, Zhicheng Jin, Jacob Blum, Shilpa Bisht, Yong Tao, Yuba Bhandari, Daniel Petkovich, Hisashi Ro, Lijing Yang, Tza-Huei Wang, Thomas R. Pisanic, Stephen B. Baylin, Hariharan Easwaran

**Affiliations:** Department of Oncology, The Sidney Kimmel Comprehensive Cancer Center at Johns Hopkins, Baltimore, MD, 21287, USA; Department of Mechanical Engineering, Johns Hopkins University, 3400 N Charles St., Baltimore, MD 21210, USA Johns Hopkins Institute for Nano BioTechnology, 3400 N Charles St., Baltimore, MD 21210, USA

## Abstract

DNA methylation alterations are hallmark features of aging and cancer, occurring early during tumor initiation. Yet how these changes evolve during early tumorigenesis, and whether oncogenic signaling actively drives them, remains poorly understood due to limitations in resolving this at the longitudinal and clonal level. Here we developed a genome-wide DNA methylation profiling method for low DNA amounts and applied it to single stem cell-derived clones from a *BRAF^V600E^* -inducible mouse colon organoid model, tracking nucleotide-level methylation heterogeneity over time. *BRAF^V600E^* induction produced stepwise methylation divergence consistent with progressive clonal takeover, in contrast to gradual epigenetic drift in uninduced controls. Strikingly, while the specific CpG sites altered differed between *BRAF^V600E^* and control lineages, they converged on the same genomic elements. Particularly, Polycomb-marked promoter CpG-island methylation gains were not induced or accelerated by *BRAF^V600E^*, challenging models in which oncogenic signaling or cell proliferation actively instructs de novo methylation of Polycomb-targeted promoters. Instead, these gains tracked with “biological aging” independent of replication, which was validated across independent aging, cancer, and cell-line datasets. *BRAF^V600E^* -induced clones additionally showed progressive loss of intestinal lineage-specification transcription factor programs, consistent with selection for cells adapted to oncogenic stress. Together, these findings show that epigenetic trajectories during early tumor development, are rooted in baseline, age-dependent drivers of epigenetic alterations, and that oncogenic signaling acts primarily by selecting and expanding clones bearing pre-existing, age-associated epigenetic drift rather than instructing new methylation. Our findings have implications for how promoter methylation is interpreted as a cancer biomarker and for identifying upstream aging-associated processes as targets for early intervention.

## Introduction

DNA methylation alterations, particularly at 5′-CpG-3′ sites in the genome, are a hallmark of aging and cancer, occurring early during tumor initiation across multiple cancer types (1–3). These alterations typically involve global or domain-specific hypomethylation with focal gains of methylation at CpG-rich promoters (CpG-island promoters) and other regulatory elements (4,5). CpG-island (CGI) promoters primarily regulated by Polycomb (PcG) group modifiers of chromatin, marked by the histone mark H3K27me3, are specifically susceptible to methylation in cancers and aging (6–10). However, the processes that determine the locus specificity and progressive accumulation of this methylation gains remain incompletely understood. Proposed, non-mutually exclusive mechanisms include oncogene-directed chromatin remodeling and DNA methyltransferase recruitment (11–14), replication-and cell-division–associated epigenetic drift (13,15), inflammation/oxidative damage (16–18), and environmental exposures (19–21). These processes constitute key factors of aging, which involves the same kind of epigenetic changes observed in cancers in apparently normal cells (10,22–25).

Colorectal cancer (CRC), particularly that driven by the *BRAF^V600E^* serrated pathway, provides a powerful model to address this question. Tumors arising from this pathway exhibit widespread promoter CpG island hypermethylation, termed the CpG Island Methylator Phenotype (CIMP) (26–28). *BRAF^V600E^* has been reported to induce DNA methylation at gene CGI promoters associated with gene silencing and has been proposed as a driver of the selected promoter CGI loci that are characteristically hypermethylated in CIMP-high CRCs (11). The proposed mechanisms range from ERK-dependent stabilization of MAFG and recruitment of DNMT3B-containing corepressor complexes (11), to methylation maintenance by Polycomb Group (PcG) components (EZH2, SUZ12, and EED) (29,30). However, other studies indicate that *BRAF^V600E^* alone is insufficient to induce methylation programs and that the methylation changes accumulate slowly or may precede, and facilitate, *BRAF*-driven transformation (22,31). The direct role of an oncogene in inducing CpG-island methylation remains incompletely tested in physiologically relevant models of early tumorigenesis that can distinguish de novo, locus-specific methylation induction.

To investigate this, we leveraged our previously established mouse organoid model of *BRAF^V600E^*-induced colon tumorigenesis as a controllable, oncogene-driven system to dissect the clonal dynamics of DNA methylation over time, processes that are difficult to resolve in vivo (22). Prior work using this model showed that *BRAF^V600E^*-induction results in organoid transformation over a period of 5-6 months in culture and is associated with DNA methylation accumulation at CGI promoter of key tumor-suppressor genes commonly hypermethylated in CIMP-H colorectal cancer, including genes involved in WNT regulation, colonic epithelial differentiation, and cell-cycle control (22). We developed a novel low-input, genome-wide whole genome bisulfite sequencing (WGBS) method for epigenetic profiling of individual organoid clones with longitudinal tracking to directly test whether oncogenic *BRAF^V600E^* instructs de novo CGI promoter methylation. We further resolve this using converging evidence from independent experimental and clinical data.

Our findings indicate that *BRAF^V600E^* does not globally induce or amplify DNA methylation gains. CGI promoter hypermethylation follows a shared, age-in-culture–associated epigenetic drift, with susceptible CpG sites converging on similar genomic elements across lineages. Notably, promoter CGI hypermethylation is uncoupled from increased cell-division cycles, whereas methylation loss at non-CGI and late-replicating regions is more consistent with replication-associated drift. Within this shared epigenetic background, *BRAF^V600E^* is associated with distinct methylation changes that are consistent with a model of selective adaptation to oncogenic stress. Together, these findings suggest that DNA methylation remodeling during tumorigenesis is not simply a direct consequence of oncogenic signaling or proliferation but instead reflects context-dependent interactions among time-dependent epigenetic drift, chromatin state, and clonal evolution.

## Results

### Low-Input WGBS Profiling of Single-Cell Derived Organoid Clones Over a 5-Month Chronological Time course

To investigate these questions, organoids were established from colonic epithelium of mice harboring a Cre-activatable *BRAF^V600E^* allele and a Rosa26-tdTomato reporter allele, enabling fluorescent labeling of cells that had undergone Cre-mediated recombination. This system was selected because our previous studies have shown that *BRAF^V600E^* induction results in phenocopying of the age-related, CIMP-H colon cancer oncogenesis (22). Following establishment, Cre recombinase was delivered via lentiviral transduction to activate *BRAF^V600E^*; parallel control lines were transduced with the lentiviral backbone without Cre-recombinase (empty vector, Ctrl). Successful recombination resulted in expression of the tdTomato-floxed marker locus (Fig. 1b). Organoids were then enzymatically dissociated into single cells monthly for five months, re-plated, and expanded into clonal organoids. The single cell disassociation efficiency was about ∼99% (Supplementary Fig. 1a). Each organoid clone is thus derived from a single stem cell. Five single organoid clones were picked at the monthly intervals and were subsequently processed for whole genome bisulfite sequencing (WGBS) (schematic in Fig. 1a).

**Figure 1:**
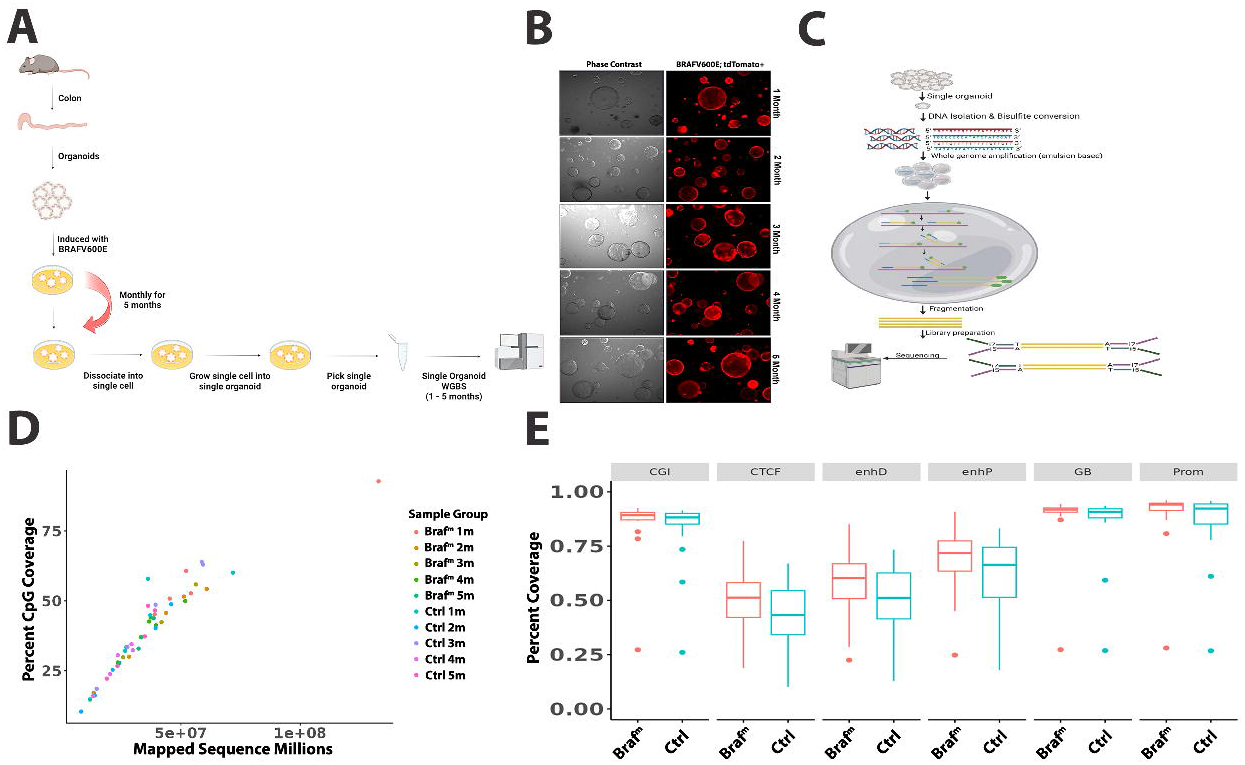
Low-Input emulsion-based WGBS Profiling of Single-Cell Derived Organoid Clones Over a 5-Month Chronological Time course. (A) Schematic of the performed investigation. (B) Representative phase-contrast (left) and *tdTomato* fluorescence (right) images of *BRAF^V600E^* expressing organoids at the indicated time points (1–5 months). (C) Schematic of droplet organoid WGBS, grey bubble indicates an emulsion based multiple displacement amplification reaction. (D) Percent CpG Coverage as a function of Mapped Sequences in Millions per lineage and timepoint. (E) Percent coverage per lineage of different genomic elements (CGI -CpG Islands, CTCF sites, enhD -Distal Enhancers, enhP -Proximal Enhancers, GB -Gene Bodies and Prom – Promoters). Enhancers defined as per the ENCODE annotation for mm10, gene bodies represented as the transcription start site to transcription end site, promoters represented as 1500bp +/-around the transcription start site.

Given the limited DNA yield from a single organoid, modifications to standard WGBS library preparation were required. We developed a simplified approach, wherein DNA from single-cell derived organoids were bisulfite converted, followed by multiple displacement amplification (MDA) in millions of picoliter-scale “oil-in-liquid” reaction chambers (emulsion droplets) using bisulfite-aware random primers (eWGBS-seq, Fig. 1c, Methods). The emulsion MDA products were purified and fragmented, and sequenced (150bp paired-end). This approach results in reads with high bisulfite conversion efficiency (>98 %, Supplementary Fig. 1b). This emulsion-based strategy reduces amplification bias and background artifacts, increasing the proportion of high-quality sequencing reads (Supplementary Fig. 1c). Across independently prepared libraries, genome-wide CpG coverage increased linearly with the number of mapped sequencing reads indicating MDA and subsequent Illumina library preparation maintained sufficient library complexity enabling consistent recovery of CpG sites with increasing sequencing depth (Fig. 1d). Evaluation of genomic features (ENCODE annotated) (32), demonstrated that CpG recovery was highest in CGIs, promoters, and gene bodies (>80%) and more moderate in CTCF-binding and enhancer-associated regions (50–70%). This pattern is consistent with greater representation of CpG-rich promoter-associated features in our low-input bisulfite-sequencing libraries, although feature-specific coverage may also reflect CpG density, mappability, genomic feature size, and library-preparation bias (33,33–35).

We evaluated the effect of CpG coverage thresholds on global methylome comparisons among individual organoid clones. We calculated pairwise Euclidean distances among organoid clones using all CpG sites covered by at least one read (≥1x; Supplementary Fig. 1d) and compared these matrices to those restricted to high-confidence sites with four or more reads (≥4x; Supplementary Fig. 1e). The resulting distance matrices exhibited a highly significant correlation (p < 0.001, Mantel test), demonstrating that single-read coverage reliably recapitulates the overall methylome architecture for independent clones, which was used for global analyses to maximize genomic coverage, including sample-level distance calculations and broad methylation-pattern comparisons.

### BRAF overexpression does not cause acute and rapid changes in DNA methylation

To define the temporal relationship between oncogenic BRAF activation and epigenetic remodeling, we performed multidimensional scaling (MDS) of genome-wide CpG methylation profiles using CpG sites shared across all single-cell-derived organoid clones. This analysis revealed distinct lineage-specific methylome trajectories over the 5-month culture period. In the *BRAF^V600E^* lineage, clones showed a more progressive, stepwise separation across successive time points, suggesting cumulative methylome alterations during long-term oncogenic evolution. In contrast, control clones exhibited a pattern consistent with epigenetic drift from months 1 to 4, followed by marked convergence at month 5 (Fig. 2a). Particularly individual clones across the 1 to 4-month time points showed varied similarity, suggesting coexistence of distinct epigenotypes within the control lineage, a feature of epigenetic heterogeneity/drift. Despite these distinct trajectories, in both control and *BRAF^V600E^* lineages we observe reduced within-lineage dispersion at 5 months, suggesting that by this time the clones undergo selection for, or convergence toward, a relatively stable epigenetic state under both oncogene-induced and -uninduced conditions.

**Figure 2:**
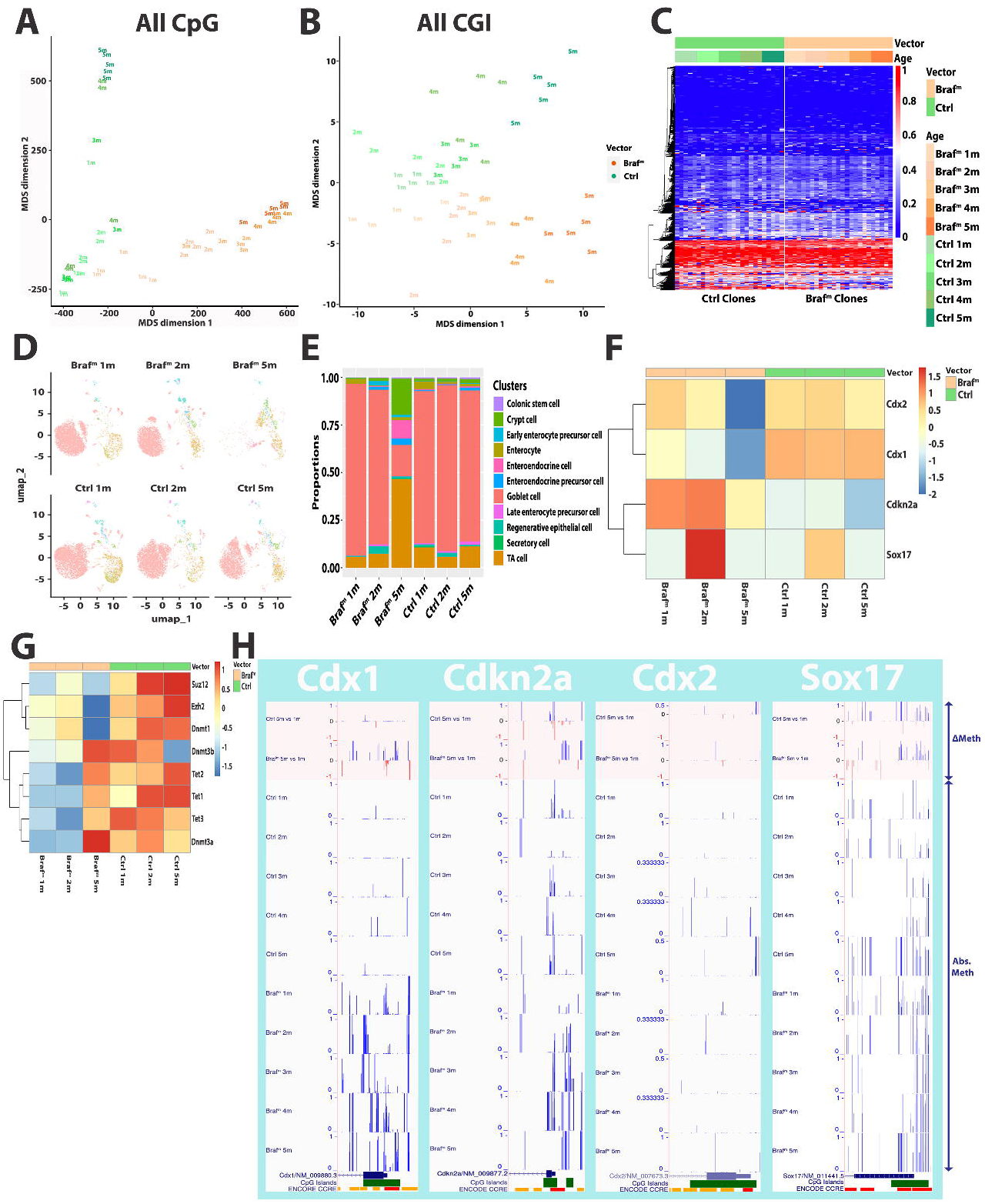
*BRAF^V600E^* overexpression does not cause acute and rapid changes in DNA methylation. (A) Multidimensional scaling of organoid WGBS across all CpG sites; organoid clones are colored by month time point and lineage (Braf^m^ – *BRAF^V600E^* lineage, Ctrl – Control lineage). (B) Multidimensional scaling of droplet organoid WGBS across all CpG Island sites, organoid clones are colored by month time point and lineage. (C) Heatmap with row based unsupervised hierarchical clustering of droplet organoid based WGBS of all common CpG sites across all clones and lineages. Heatmap annotations represent organoid clone month timepoints and lineage. (D) UMAP of scRNA-seq of organoid clones at 1-month, 2-month and 5-month timepoints split by lineage. (E) Proportions of annotated cell types represented in each month timepoint across lineages. (F) Heatmap of pseudobulk scRNA-seq of common colon cancer related genes with row based unsupervised hierarchical clustering across all clones and lineages. (G) Heatmap of pseudobulk scRNA-seq of common mechanistic components relating to the deposition and removal of DNA methylation across month timepoints and lineages. (H) UCSC Genome Browser tracks showing the difference in methylation (ΔMeth -orange shaded boxes) and absolute methylation for each timepoint and lineage across the same colon cancer related genes shown in F.

At single-CpG resolution, genome-wide CpG methylation profiles indicated that *BRAF^V600E^* induction did not result in an immediate, global shift in DNA methylation distributions (Fig. 2c). CpG methylation levels were overall similar between uninduced control and *BRAF^V600E^* clones during the early months of culture. This similarity was maintained when analyses were restricted to all CGIs, Polycomb repressive complex 2 (PRC2)-associated CGIs, or non-CGI intergenic regions (Supplementary Fig. 2a–c). This observation contrasts with prior reports suggesting that *BRAF^V600E^* induces acute and widespread CGI promoter DNA methylation gains (11). The overall similarity in global methylation levels, together with the progressive separation of *BRAF^V600E^* and control clones in MDS space, suggests that oncogenic-BRAF may initially alter the trajectory, variance, and clonal selection of epigenetic states rather than directly inducing a large, uniform shift in CpG methylation. Over time, the altered evolutionary landscape may permit the accumulation or selection of locus-specific methylation changes associated with transformation.

### Gradual remodeling of cellular programs and gene expression patterns

To determine changes in cellular programs during this timeframe, we performed single-cell RNA sequencing (scRNA-seq) on the control and *BRAF^V600E^*-induced organoids at 1-, 2-, and 5-month time points. In agreement with the methylome profiling, oncogene activation resulted in a gradual remodeling of the epithelial lineage rather than acute cell state transformation. While control organoids maintain overall cell-type composition through 5 months of culture, the *BRAF^V600E^* lineage exhibited a profound, progressive loss of goblet cells starting at month-1 and culminating by month-5 (Fig. 2d). This loss was accompanied by accumulation of transit-amplifying cells, enterocytes/enteroendocrine cells, and crypt cells (Fig. 2e). Importantly, these prominent lineage shifts only fully emerged at the 5-month time point, indicating that the phenotypic features of *BRAF^V600E^*-driven dysplasia evolve across successive cell generations rather than appearing as an immediate post-induction cellular response.

In response to *BRAF^V600E^* induction, secretory, transit-amplifying, and crypt-like populations displayed robust enrichment of MAPK/ERK transcriptional signatures following *BRAF^V600E^* induction, whereas colonic stem cells showed comparatively attenuated MAPK/ERK transcriptional output (Supplementary Fig. 2d). Excessive or sustained MAPK/ERK activation can promote loss of canonical intestinal stem-cell identity through differentiation and/or senescence-like responses and can activate negative-feedback regulators, including DUSP4, DUSP5, DUSP6, and SPRY4 (Supplemental Fig. 2d, Erk Feedback Regulators) (36–39). The relatively attenuated MAPK/ERK signature in colonic stem cells may reflect cell-state-specific modulation of the response to BRAF^V600E^, consistent with reports that reduced ERK phosphorylation can enhance intestinal stemness in BRAF^V600E^-expressing epithelium and that stem/TA-like colorectal cancer cells exhibit relatively low MAPK-target expression (36,40). Such modulation could favor maintenance of a stem-like state during sustained oncogenic signaling. In contrast, stronger MAPK/ERK signaling in secretory and transit-amplifying populations may contribute to altered differentiation dynamics and reduced goblet-cell lineage.

To gain insights into the transcriptional control mechanisms governing the lineage destabilization and early steps of transformation, we conducted pseudobulk scRNA-seq analysis of key colorectal cancer-associated transcription factors (TFs) and tumor suppressors (Fig. 2f) (31,41–46), whose downregulation was shown to be important for *BRAF^V600E^*-induced tumorigenesis (22,47). We observed progressive and robust downregulation of the homeodomain transcription factors Cdx1 and Cdx2. Loss of CDX2 is a clinical hallmark of the *BRAF^V600E^* serrated pathway, where epigenetic silencing of CDX2 and subsequent protein loss are tightly associated with CIMP-H status, mismatch repair deficiency, right-sided tumor location, and aggressive histomorphological features (42,48,49). In our previous work developing these organoid models, CDX2 downregulation is a critical molecular driver of lineage dedifferentiation and its loss primes *BRAF^V600E^*-induced transformation (22,49).

*BRAF^V600E^* induction resulted in a marked upregulation of the key tumor suppressor and senescence mediator *Cdkn2a* (p16) (31,43) at months 1 and 2, with relative suppression by month-5 (Fig. 2f). This temporal pattern reflects initial oncogene-induced senescence, a protective barrier against hyperactivated MAPK signaling, followed by epigenetic escape and Cdkn2a downregulation to drive clonal expansion (31,43). The developmental transcription factor *Sox17* (50) exhibited a sharp increase in the second month post-induction before returning to baseline levels, indicating early, dynamic changes in cell-state or lineage-specification response rather than an immediate and sustained change in cellular identity after *BRAF^V600E^* induction (Fig. 2f).

Given that genes such as *CDKN2A*, *SOX17*, *CDX2* are frequently hypermethylated in BRAF mutant, CIMP-H CRCs, we investigated the promoter CGI-methylation at these loci, and other genes frequently hypermethylated in CIMP-H CRCs. Longitudinal methylation analyses revealed distinct gene-specific trajectories, indicating that the methylation changes are heterogeneous, gradual and show locus-specific temporal patterns. *Cdkn2a* showed a gradual increase in CGI methylation beginning 2–3 months after BRAF induction, closely following the *Cdkn2a* gene repression pattern (Fig. 2h). Control organoids show substantially smaller and heterogenous methylation gains in *Cdkn2a*. *Cdx1* showed sustained methylation gain from the earliest time point measured, aligning closely with its early, progressive transcriptional downregulation. In contrast, *Cdx2* showed no appreciable methylation gain over the time course, although it undergoes transcriptional repression by 5-months post-BRAF induction, indicating that within this window *Cdx2* silencing is potentially regulated by other repressive mechanisms. This paralog-specific divergence between Cdx1 and Cdx2 illustrates that even closely related, co-regulated transcription factors can be silenced through distinct chromatin mechanisms during early transformation. Sox17 exhibited a similar progressive methylation-gain trajectory in both control and *BRAF^V600E^* lineages, indicating a shared time-dependent, rather than oncogene-dependent, process. Similar patterns were observed for Sfrp2 and Sfrp4 (Supplemental Fig. 2e). These gene-specific trajectories indicate that *BRAF^V600E^* does not uniformly induce promoter CGI hypermethylation. Instead, methylation changes in the BRAF-induced lineage emerge gradually at a subset of loci, while other promoter CGIs follow lineage-independent, culture-associated methylation trajectories.

To explore roles of epigenetic modulators in the observed methylation trajectories, we examined the expression of key DNA methylation and chromatin-regulatory factors, including DNA methyltransferases (*Dnmt1*, *Dnmt3a*, *Dnmt3b*), TET dioxygenases (Tet1–3), and PRC2 components (Fig. 2g). Expression of these factors was generally lower in the *BRAF^V600E^* lineage than in controls at the early 1-and 2-month time points; *Suz12*, *Ezh2*, and *Dnmt1* remained lower throughout the 1-to 5-month time course, whereas *Dnmt3a*, *Dnmt3b*, and *Tet1-3* starkly increased by 5 months. Expression pattern of these key methylation-regulatory factors argues against a model in which oncogenic-BRAF signaling drives a sustained, uniform increase in DNA methyltransferase or Polycomb-factor expression; instead, it suggests an early restriction in gene expression. The locus-specific roles of these factors remain to be addressed.

Together, these results demonstrate that *BRAF^V600E^* overexpression is not associated with acute, genome-wide de novo DNA methylation remodeling. Similarly, at the transcriptional level, *BRAF*^V600E^ triggers the expected early gene response to oncogenic stress, with eventual progressive lineage destabilization and cellular stress responses. The data suggests locus-specific epigenetic changes, and a transcriptional landscape progressing towards increased proliferation state, over successive cell generations during the early steps of *BRAF^V600E^*-induced transformation.

### Genomic context, not oncogenic BRAF, determines the direction of methylation change

To resolve the precise genomic sites subject to DNA methylation in the *BRAF^V600E^* and control lineages over the 5 months culture, we performed differential methylation analysis comparing pooled 1m and 5m clones within both the control and *BRAF^V600E^* lineages (FDR < 0.05 and absolute Δ Methylation Level > 0.2). This CpG-site level temporal analysis revealed that both wild-type and oncogene-expressing lineages exhibit remarkably similar overall numbers of DNA methylation alterations, with nearly equivalent numbers of CpG sites undergoing significant methylation gains over the five-month time course; losses being more prominent in the *BRAF^V600E^*-lineage (Fig. 3a). Strikingly, despite these parallel global kinetics, the specific single-nucleotide CpG sites undergoing methylation changes showed minimal overlap between the two lineages (Fig. 3b; red = gains, blue = losses; Supplementary Fig. 3d: non-promoter CGI; Supplementary Fig. 3e: non-promoter, non-CGI).

**Figure 3:**
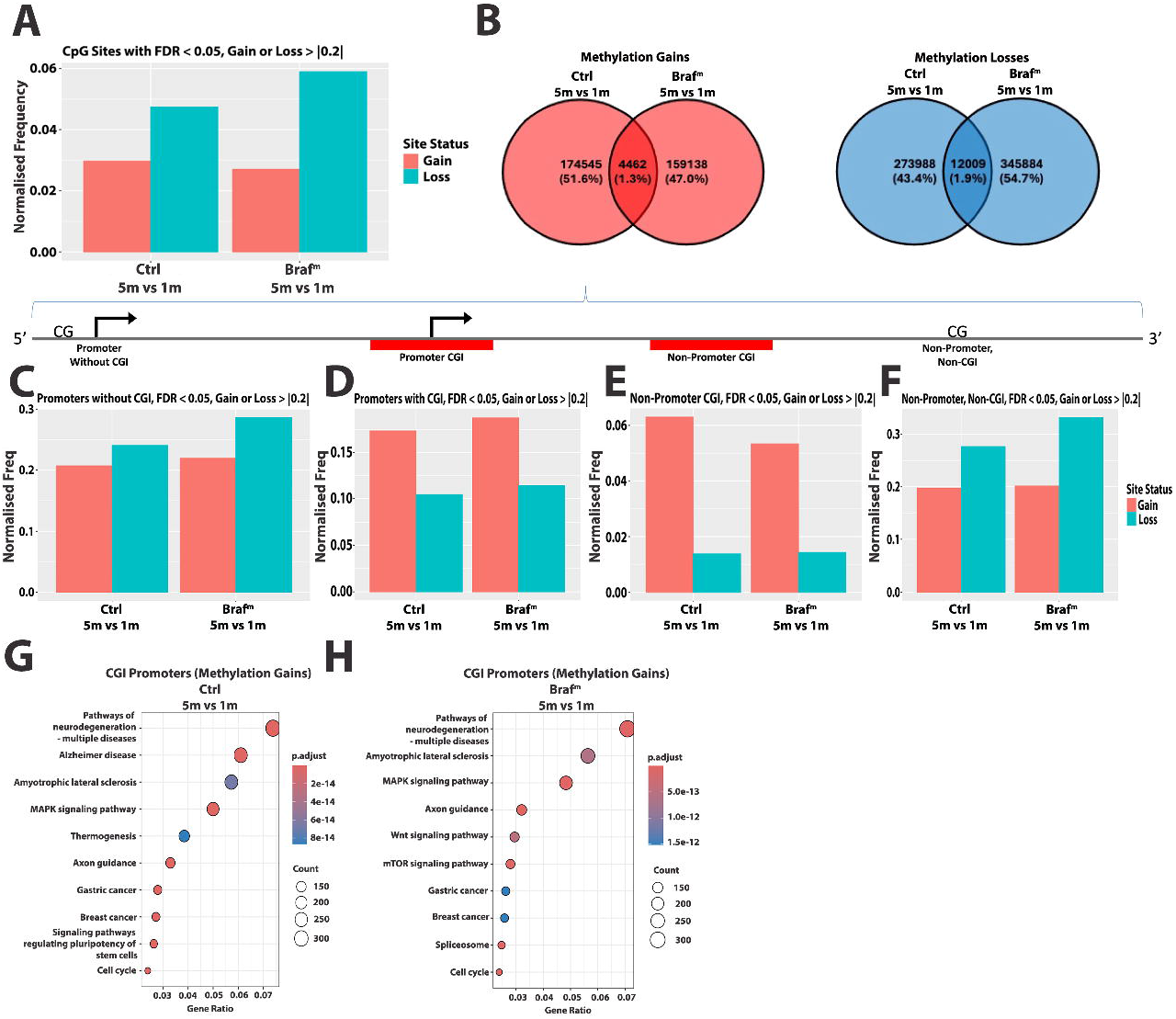
Genomic context, not oncogenic BRAF, determines the direction of methylation change. (A) Normalized frequency of differentially methylated CpG sites in 1-month vs 5-month comparison for each lineage (red represents hypermethylated CpG sites, blue represents hypomethylated sites relative to the indicated comparison). (B) Venn diagram representing the shared hypermethylated (red) and hypomethylated (blue) CpG sites between 1-month and 5-month comparisons across lineages. (C-F) Normalized frequency of differentially methylated CpG sites in 1-month vs 5-month comparison for each lineage (red represents hypermethylated CpG sites, blue represents hypomethylated sites relative to the indicated comparison) across different genomic elements; Promoters without CGI (C), Promoters with CGI (D), Non-Promoter CGI (E), Non-Promoter/Non-CGI (F). (G) KEGG over representation analysis of differentially hypermethylated sites in promoters with CGI within control (Ctrl) 1-month vs control 5-month. (H) KEGG over representation analysis of differentially hypermethylated sites in promoters with CGI within BRAF^V600E^ (Braf^m^) 1-month vs *BRAF^V600E^* 5-month. To account for differences in the number of evaluable CpG sites across time-point and lineage comparisons, the numbers of CpG sites exhibiting significant methylation gains or losses within each genomic category were divided by the total number of evaluable CpG sites in the corresponding comparison. The resulting proportions were reported as normalized frequencies.

To determine underlying genomic features of the sites with these methylation alterations, we stratified the differentially methylated CpGs by their genomic feature annotations (promoters ± CGIs, non-promoter CGIs, and non-promoter non-CGIs; Figs 3c-f). Regions containing CGIs predominantly gained methylation in both the control and *BRAF^V600E^* lineages (Fig. 3d, e), whereas regions lacking CGIs (non-CGI promoters and intergenic heterochromatic domains) predominantly lost methylation (Fig. 3c, f). This pattern of methylation changes recapitulates the well-documented dual-epigenetic hallmarks of mammalian tissue aging, chronological senescence, and early tumor initiation (51,52). Importantly, *BRAF^V600E^* induction was not observed to result in a pronounced increase in CGI methylation over control in the 5-month duration (Fig. 3d, Supplementary Fig. 2A).

KEGG pathway enrichment analysis of genes linked to differentially methylated CGI-containing promoter regions identified both shared and *BRAF^V600E^*-specific epigenetic programs in the uninduced control and mutant lineages (Fig. 3g, h; hypomethylated regions are shown in Supplementary Fig. 3b). Shared pathways involve conserved developmental and cellular homeostatic networks, including pathways of neurodegeneration, amyotrophic lateral sclerosis, MAPK signaling, axon guidance, cell cycle control, and signatures associated with gastric and breast cancer. These broad disease-associated annotations likely reflect convergence on shared processes, such as signal transduction, cytoskeletal organization, mitochondrial function, proteostasis, and RNA metabolism, rather than acquisition of disease- or tissue-specific phenotypes. The substantial overlap between control and mutant lineages suggests that culture-associated epigenetic drift contributes to remodeling of these pathways independently of oncogenic-BRAF signaling.

Pathways associated with CGI hypermethylation uniquely in *BRAF^V600E^* mutant clones include WNT signaling, mTOR signaling, and spliceosome-associated genes (Fig. 3g, h). This suggests that within a background of baseline epigenetic drift that affects shared pathways, the *BRAF^V600E^* oncogenic context modulates epigenetic remodeling toward pathways that regulate intestinal stem-cell identity, growth and metabolic control, and RNA processing.

### *BRAF^V600E^*-associated methylation changes converge on intestinal lineage-specification transcription factors

As noted previously, the two lineages (*BRAF^V600E^* and Control) share few differentially methylated sites (Fig. 3b, Supplementary Fig. 3d, e). However, when these sites are annotated by genomic context (e.g., promoter with CGI), there is substantial overlap between lineages (Fig. 4a), as is the pathway analyses. This pattern is consistent with shared constraints on the genomic regions susceptible to methylation remodeling during long-term culture, while the specific CpG sites affected may be lineage dependent.

**Figure 4:**
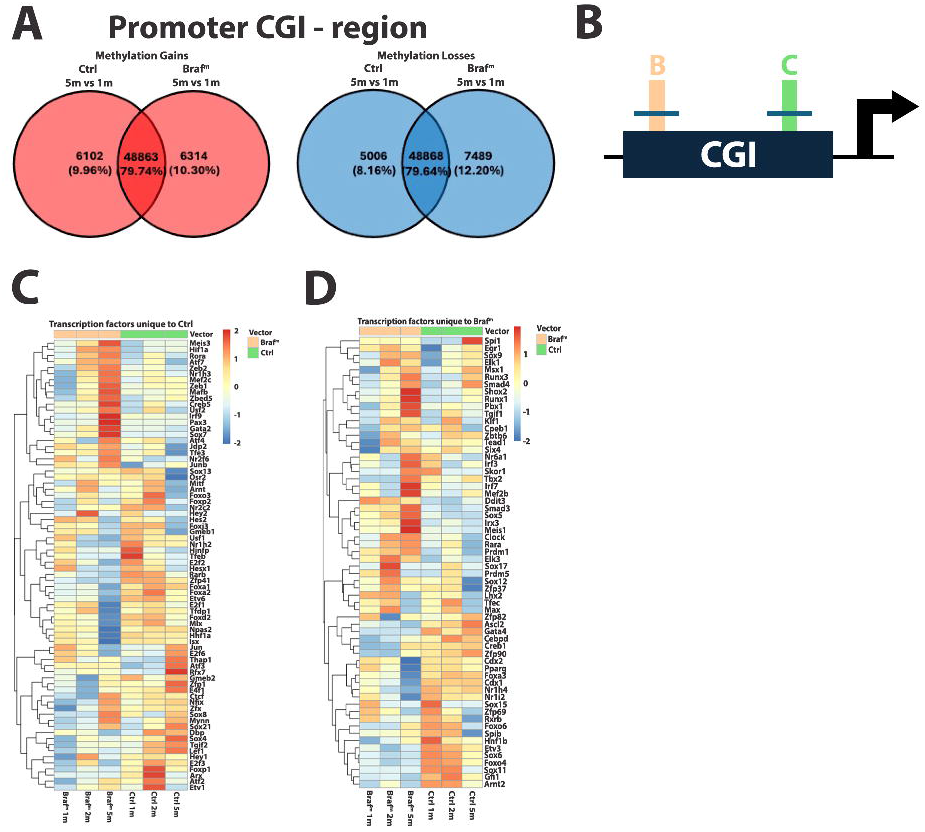
Transcription factor analyses of CGI regions with methylation changes. (A) Venn diagrams showing the overlap of differentially methylated promoter CGI regions between control and *BRAF^V600E^* organoids. (B) Schematic of the transcription factor binding enrichment analysis using differentially methylated CpG sites within promoter CGIs from BRAF^V600E^ (orange, B) and control (green, C) organoids. Blue horizontal lines indicate the 50-bp flanks added to each differentially methylated site to define the regions used for enrichment analysis. (C, D) Heatmaps of pseudobulk scRNA-seq expression at months 1, 2, and 5 in both organoid lineages for transcription factors uniquely enriched in control (H) or *BRAF^V600E^* (I) organoids. Transcription factors were identified by simple enrichment analysis (SEA from MEME Suite) of differentially methylated promoter CGI sites with 50-bp flanking sequences. Rows were ordered by unsupervised hierarchical clustering.

Previous studies have shown that methylation changes at specific genomic sites are correlated with changes in transcription factor activities (53,54). We thus explored the relationship between lineage-specific differential methylation at CGI promoters and candidate transcriptional regulators, by identifying the transcription factors whose binding motifs were enriched among differentially methylated promoter CGI sites and quantifying their expression from pseudobulk scRNA-seq data across the 1-, 2-, and 5-month time points. To this end, differentially methylated CpG sites within promoter CGIs were extended by 50 bp flanking regions, and non-overlapping, lineage-specific regions were analyzed using Simple Enrichment Analysis (MEME Suite) to identify enrichment of transcription factor binding motifs (Fig. 4b).

*BRAF^V600E^*-associated differentially methylated CpG sites are enriched for motifs of TFs that converge on intestinal epithelial cell-state transition and factors important in colorectal cancers, such as crypt/stemness and WNT-associated regulation (SOX9, ASCL2, SOX17, CDX1/2), intestinal lineage specification (CDX1/2, GATA4, FOXA3, HNF1B, NR1H4/FXR, and NR1I2/PXR) (Fig. 4d). scRNA-seq pseudobulk data revealed that of these enriched TF motifs, those of important intestinal lineage regulators specifically Cdx1, Cdx2, Foxa3, Pparg, Nr1h4/Fxr and Nr1i2/Pxr are stably expressed across all time points in uninduced control organoids, while these factors are downregulated by 5m in the *BRAF^V600E^* lineage (Fig. 4d).

Conversely, a subset of transcription factors with motifs enriched in the *BRAF^V600E^* lineage exhibited progressive transcriptional upregulation over the 5-month time course (Fig. 4d). This cluster included regulators of intestinal progenitor-state plasticity and developmental transcriptional programs (SOX9, TEAD1, PBX1, MEIS1, and SIX4); ERK-responsive immediate-early transcription (EGR1, ELK1, and ELK3); and modulators of TGF-β/SMAD signaling (SMAD3, SMAD4, and TGIF1). The cluster also contained factors associated with integrated cellular stress and innate inflammatory responses, such as DDIT3, IRF3, and IRF7. Together, these changes are consistent with progressive BRAF-associated remodeling of epithelial cell-state regulatory networks, integrating mitogenic signaling, stress adaptation (31,43), and regenerative or fetal-like plasticity (37,55). This state was most pronounced at 5 months, suggesting that sustained BRAF^V600E^ signaling promotes progressive lineage destabilization and adaptation.

The control lineage’s differentially methylated CpG sites were enriched for motifs broadly linked to stress and antiviral responses (Fig. 4c), involving key components of the AP-1 stress-response complex (JUNB, JUN, ATF3), cellular senescence regulators (FOXO3), and mediators of antiviral interferon and inflammatory signaling (IRF9, NFKB1, NFKB2) (56–60). Pseudobulk scRNA-seq demonstrated that these stress and immune-surveillance TFs are selectively upregulated in uninduced control clones over time, highest levels at month-5 (Fig. 4c). The enrichment of stress-response TF-motifs, together with the corresponding temporal expression patterns, suggest engagement of stress-adaptation regulatory programs in uninduced organoid clones during chronological aging in culture.

The above data support a model in which both lineages undergo a similar degree of time-dependent, culture-associated CGI-methylation changes. However, oncogenic *BRAF^V600E^* reshapes the landscape of this process, but does not acutely induce methylation changes. In the *BRAF^V600E^* lineage, progressive silencing of intestinal differentiation regulators, including Cdx1 and Cdx2, is consistent with erosion of the transcriptional programs that maintain mature colonic epithelial identity, promoting transition toward a less differentiated, dysplastic cell state. Concurrent enrichment of stem-and plasticity-associated regulators, including Sox9, together with altered Runx3-, Smad4-, and other differentiation-associated regulatory networks, suggests selection for cell states that can tolerate oncogenic-*BRAF* signaling, evade oncogene-induced senescence (22,37,49).

### DNA Methylation gains at promoter CGI independent of *BRAF^V600E^* induction

We characterized in detail the methylation dynamics over time to distinguish the methylation trajectories between *BRAF^V600E^* and control lineage. To this end, we identified CpG sites that showed substantial methylation changes between 1 and 5 months (Δ methylation > 0.5 for gains and Δ methylation < -0.5 for losses). We then tracked methylation levels at these sites across all intermediate timepoints in each lineage and genomic context (schematic in Fig. 5a).

**Figure 5:**
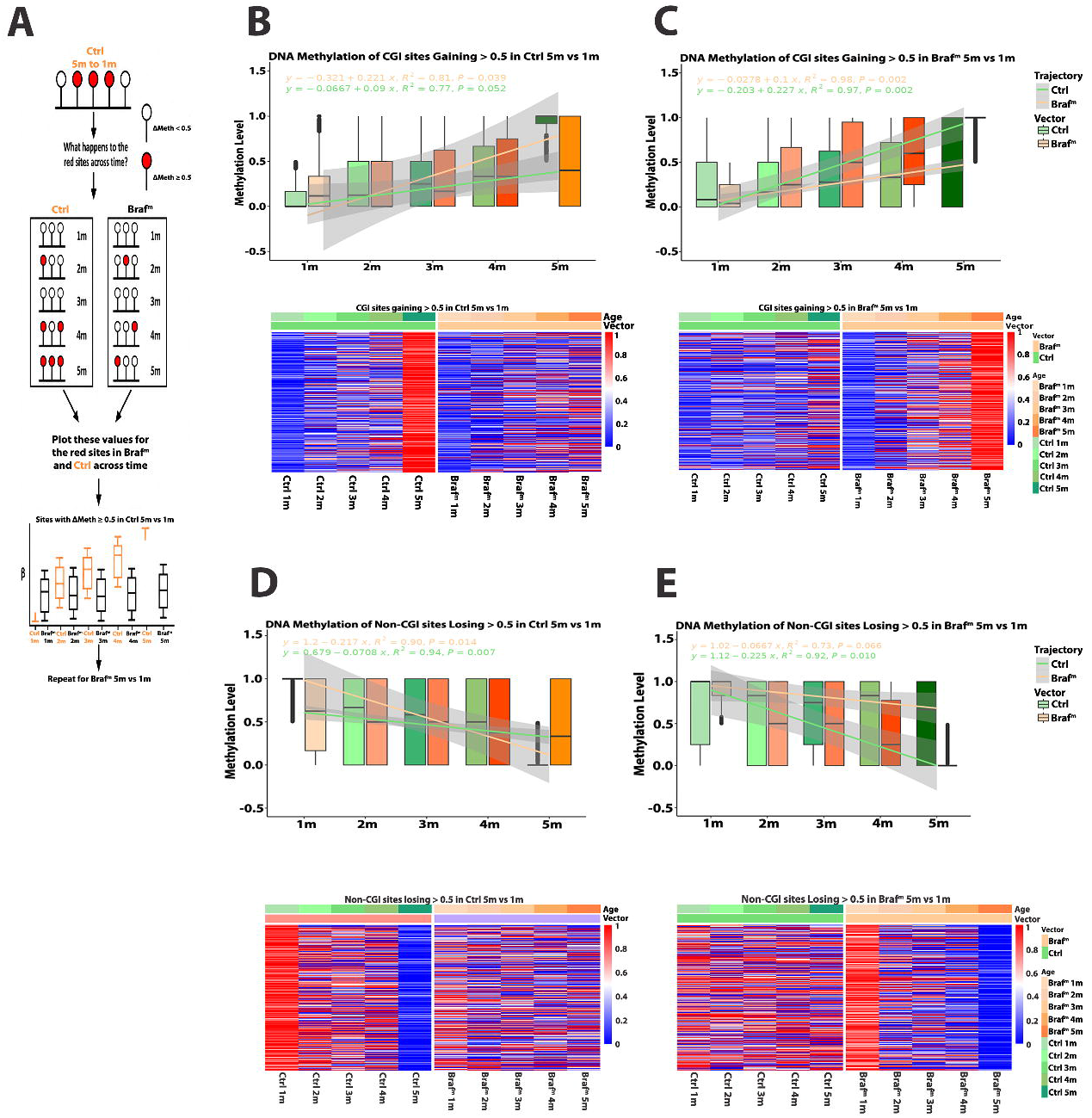
DNA Methylation gains occur at promoter CGI independent of *BRAF^V600E^* induction. (A) Longitudinal analysis of DNA methylation at CpG sites exhibiting increased methylation between months 1 and 5. Sites with a methylation increase of at least 0.5 (ΔMeth ≥ 0.5) in control organoid between month 1 and month 5 are selected, and methylation β values at these same sites are examined in both control (Ctrl) and BRAF-mutant (Braf) cultures at each month time point. Red and unfilled circles indicate sites that meet or do not meet the selection threshold, respectively. The schematic boxplots is a representative illustration of distributions of methylation values across the selected sites in each condition over time (Ctrl, orange; Braf, black). The analysis is then repeated using sites selected by the same criterion between months 1 and 5 in Braf organoids. (B) Longitudinal analysis of DNA methylation at CpG sites within CpG Islands exhibiting increased methylation between months 1 and 5 of the control lineage and the representative methylation of those same sites in the *BRAF^V600E^* lineage. (C) Longitudinal analysis of DNA methylation at CpG sites within CpG Islands exhibiting increased methylation (ΔMeth ≥ 0.5) between months 1 and 5 of the *BRAF^V600E^* lineage and the representative methylation of those same sites in the control lineage. (D) Longitudinal analysis of DNA methylation within non-CpG Island CpG sites exhibiting decreased methylation (ΔMeth ≤ 0.5) between months 1 and 5 of the control lineage and the representative methylation of those same sites in the *BRAF^V600E^* lineage. (E) Longitudinal analysis of DNA methylation within non-CpG Island CpG sites exhibiting decreased methylation (ΔMeth ≤ 0.5) between months 1 and 5 of the *BRAF^V600E^* lineage and the representative methylation of those same sites in the control lineage. Heatmap below each boxplot shows the methylation of all selected non-CpG Island related CpG sites across lineages and time. Selection criteria of the sites investigated can be observed in the plot title.

Within CGI-associated regions, sites that gained methylation (Δ methylation > 0.5) in the control lineage also showed progressive methylation gains at the corresponding sites in the *BRAF^V600E^* lineage, and vice versa (Fig. 5b-c). For sites identified in the control lineage, methylation increased with an estimated slope of approximately 20% per month in control organoids (p=0.04) and 10% per month in *BRAF^V600E^* organoids (p=0.05). Conversely, sites identified as gaining methylation in the *BRAF^V600E^* lineage also increased in methylation in control organoids, with estimated slopes of approximately 20% per month in the mutant lineage and 10% per month in the control lineage (p=0.002; Fig. 5c). Thus, promoter CGI sites susceptible to methylation gain were largely shared between lineages and followed similar time-dependent trajectories. This pattern is consistent with a basal, culture-associated tendency for promoter CGI hypermethylation that is not uniquely dependent on *BRAF^V600E^* induction. In contrast, the few CGI sites undergoing methylation loss (Δ methylation < -0.5) were not concordant between the control and *BRAF^V600E^* lineages, indicating that losses in the CGI-regions are less conserved (Supplementary Fig. 4a, b).

A distinct pattern was observed in non-CGI regions. Here, methylation losses were conserved between lineages (Fig. 5d, e), whereas methylation gains showed limited cross-lineage concordance (Supplementary Fig. 4c, d). Thus, the methylation dynamics are strongly dependent on genomic context: promoter CGI regions preferentially undergo shared time-dependent methylation gains, whereas non-CGI regions exhibit shared methylation losses. The direction of these changes reflects intrinsic differences in the susceptibility of CGI and non-CGI regions to culture-associated epigenetic drift (61–64).

### Methylation gains in CGI are replication independent

The methylation trajectory analysis above indicated that CGI-associated methylation gains occur in both lineages at similar rates, suggesting that this component of the methylation program is primarily associated with time-dependent, culture-related processes rather than being uniquely induced by *BRAF^V600E^*. Previous studies have linked promoter CGI hypermethylation and non-CGI hypomethylation to cumulative mitotic history and replication-associated remodeling of the DNA methylome (47,62–67). Particularly, age-associated methylation gains are frequent at the promoter-associated and Polycomb-regulated CGI (PcG-CGI), whereas methylation loss is more characteristic of non-CGI and late-replicating genomic domains (63,68–71). Because *BRAF^V600E^* activates MAPK signaling and can increase proliferation in intestinal epithelial cells, it may increase the number of replication cycles over which such methylation alterations can accumulate. We therefore examined the distribution of differentially methylated sites across replication-timing domains (72). Consistent with established patterns of replication-associated epigenome remodeling, methylation losses were preferentially enriched in late-replicating regions, and this enrichment was greater in the *BRAF^V600E^* lineage than in controls (Fig. 6a). These patterns are potentially due to the increased proliferative history in *BRAF^V600E^* lineage that contributes to replication-associated methylation loss, prominent in late-replicating genomic domains.

**Figure 6:**
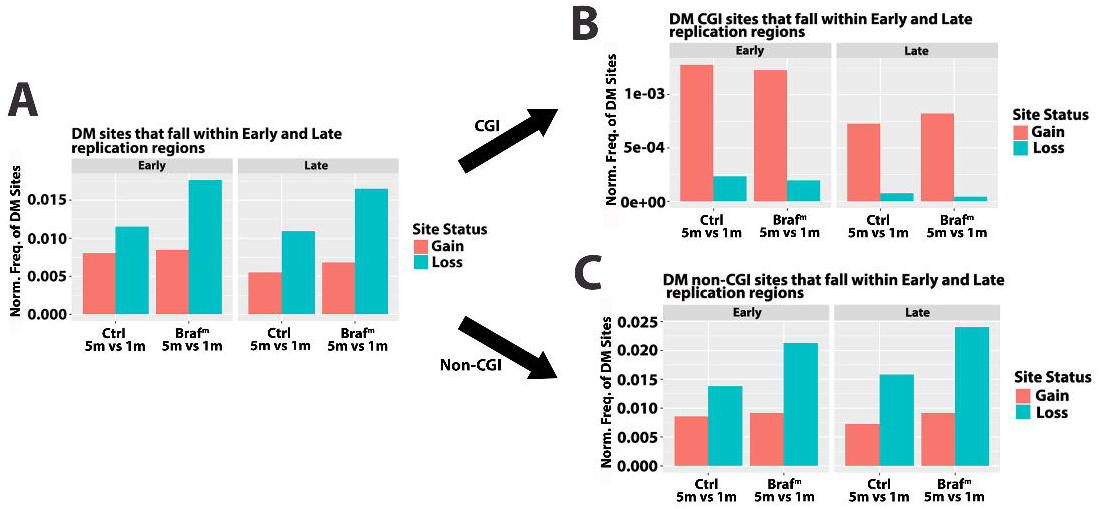
Methylation gains in CGI are decoupled from replication. (A) Normalized frequency (Norm. Freq.) of Hypermethylated (red) and hypomethylated (blue) differentially methylated (DM) CpG sites which reside within early or late replication regions across control (Ctrl) and *BRAF^V600E^*(Braf^m^) 1 – 5month comparisons. Black arrows indicate similar analysis but for differing genomic elements; (B) CGI, (C) Non-CGI.

Stratification by genomic context revealed distinct patterns. Methylation gains within CGI-associated regions were highly similar between control and *BRAF^V600E^* lineages across replication-timing domains i.e. early or late replication regions as delineated in (72) (Fig. 6b), indicating that these gains occur independently of both *BRAF^V600E^* induction and replication history. In contrast, non-CGI regions showed more prominent hypomethylation in the *BRAF^V600E^* lineage, particularly within late-replicating domains (Fig. 6c). Thus, the BRAF-associated difference was driven primarily by genomic context-dependent methylation loss rather than by a generalized increase in CGI methylation.

Based on the above findings, we focused on methylation trajectories in PcG-marked CGI regions. These regions displayed highly similar hypermethylation-rate trajectories in both lineages (Fig. 7a), further supporting the interpretation that CGI hypermethylation represents a baseline, time-dependent process rather than a specific consequence of oncogenic-BRAF signaling. In contrast, non-CGI regions showed the expected replication-associated methylation loss, with a steeper hypomethylation rate observed in the *BRAF^V600E^* lineage (Fig. 7b).

**Figure 7:**
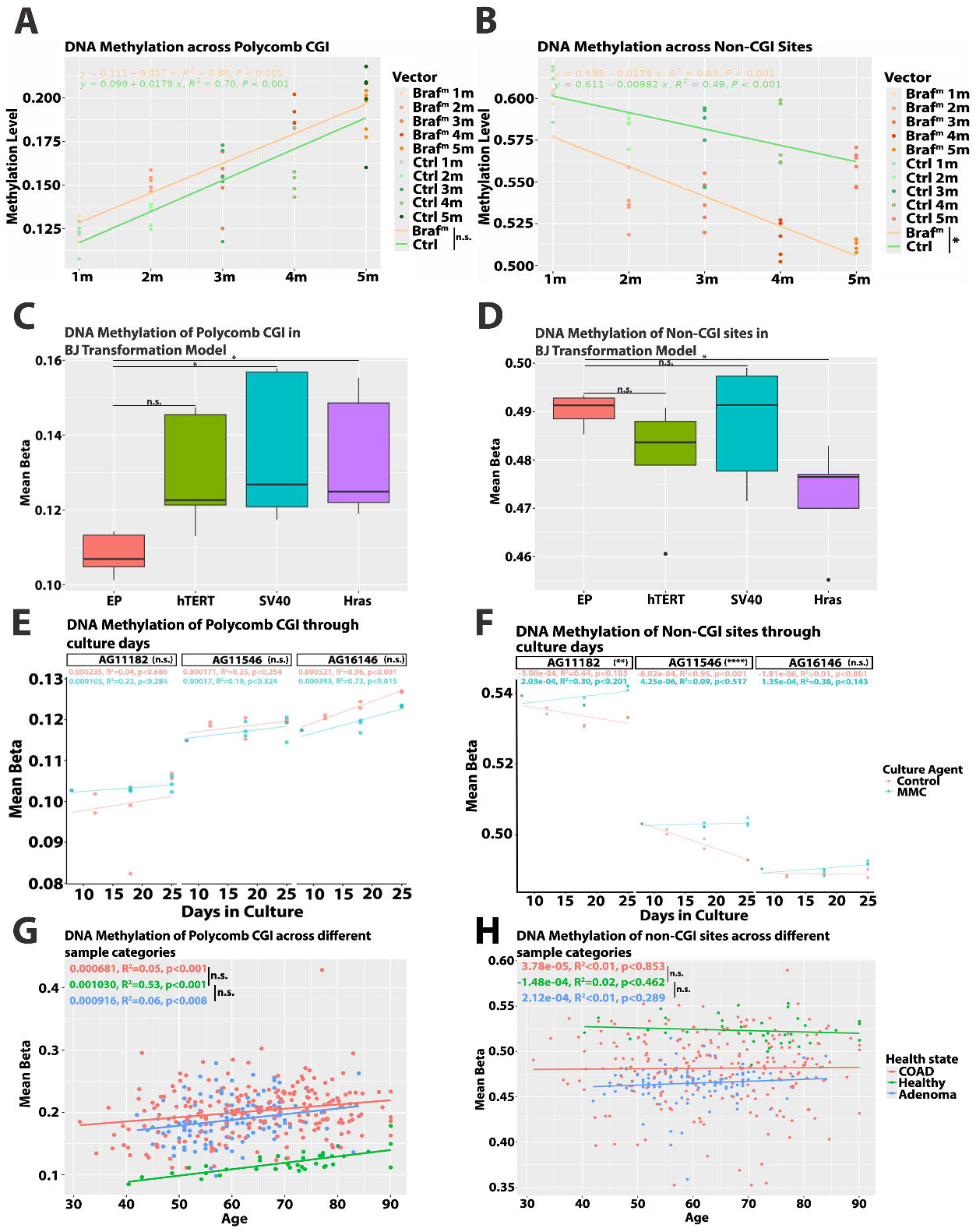
Methylation gains in promoter CGI are replication independent. (A, B) Longitudinal analysis of the DNA methylation at Polycomb marked CGI (A) or Non-CGI sites (B) across control (Ctrl) and *BRAF^V600E^* (Braf^m^) lineages and month timepoints. Individual organoid clones colored by lineage (orange – *BRAF^V600E^*, green – Control) and timepoint, lines represent a smoothed linear model with equations, R^2^ and p values noted per lineage. P-values relating to interactions between models are shown via a vertical line listed in the graph legend (A, Control vs *BRAF^V600E^* models not significantly different; B, Control vs *BRAF^V600E^* models are significantly different). (C) Investigation of the DNA methylation of polycomb marked CGI or non-CGI (D) sites upon sequential addition of proliferative agents, data from GSE91069. (E) Investigation of DNA methylation of polycomb marked CGI or non-CGI (F) sites across culture days within different cell lines treated with Mitomycin C (MMC) or control treatments. Cell line identifiers noted in the graph facet title, interaction p-value noted in brackets within the facet title, linear model slope values, R^2^ and associated linear model p-value colored per intervention. (G) Investigation of DNA methylation in polycomb marked CGI or non-CGI (H) sites in TGCA COAD and tubular adenoma from GSE233616. Lines represent linear models with slope values, R^2^ and p-values colored per health state. Vertical lines between equations indicate interaction p-values between linear models. N.s. (p>0.05), * (≤0.05), ** (≤0.01), *** (≤0.001), **** (≤0.0001). Wilcox signed rank test unless otherwise specified. Abbreviations: EP – Early Passage, COAD – Colon adenocarcinoma, MMC – mitomycin C

### Orthogonal validation distinguishes CGI methylation gain from replication-associated hypomethylation

Above observations contrast with previous studies that have linked CGI methylation to proliferative cycles and oncogenic signaling (11,47,65–67). Therefore, we examined several independent datasets to robustly assess whether Polycomb-associated CGI methylation gain is coupled to proliferative expansion in the same manner as non-CGI methylation loss.

First, we analyzed an independent Illumina HumanMethylation450 dataset generated using the classical BJ fibroblast transformation model (GSE91069) (73). In this system, BJ fibroblasts were sequentially transduced with hTERT, SV40 early-region proteins, and oncogenic H-RAS^V12^. Following immortalization and inactivation of the p53/RB-dependent checkpoints by hTERT and SV40 early-region proteins, introduction of oncogenic H-RAS^V12^ promotes increased, sustained proliferation, and transformation, enabling anchorage-independent and tumorigenic growth (73–75). The lineages were maintained in culture for same duration during the sequential transduction process (73).

If Polycomb-CGI methylation gains were primarily determined by the number of replication cycles or by the proliferative state associated with transformation, progressive methylation increases would be expected across the transformation series, particularly in the H-RAS lineage. Examination of the methylation levels at Polycomb-marked CGI revealed that these sites remained relatively stable across the *hTERT*, *hTERT+SV40*, and *hTERT+SV40+H-RAS* transduced lineages (Fig.76c). In contrast, non-CGI regions exhibited significantly more methylation loss after *H-RAS* introduction (Fig. 7d), consistent with replication-associated hypomethylation in non-CGI and late-replicating regions. These independent data therefore support the model in which CGI methylation gains are not a simple consequence of increased oncogenic signaling, transformation or proliferative expansion.

We verified these results with methylation dynamics at solo-WCGW CpG sites, which are sites within partially methylated domains that have been shown to undergo robust, replication-associated methylation loss (62,63). In our organoid dataset, solo-WCGW sites exhibited significantly more negative methylation trajectory in the *BRAF^V600E^* lineage than in controls (Supplementary Fig. 5a), indicating methylation loss with the increased replication cycles in the *BRAF^V600E^*-induced organoids. A similar pattern was observed in the BJ transformation dataset, in which the fully transformed H-RAS-expressing state was associated with greater solo-WCGW hypomethylation (Supplementary Fig. 5b). These findings are consistent with the progressive hypomethylation at replication-sensitive non-CGI sites during increased mitotic cycles under oncogenic-signaling.

To directly test the relationship between cell division and methylation dynamics, we analyzed published primary-cell datasets in which cells were cultured under proliferative conditions or in the presence of mitomycin C, which inhibits DNA synthesis and cell division. Under proliferative conditions, solo-WCGW sites showed progressive methylation loss, whereas this loss was ablated when cell division was inhibited (Supplemental Fig. 5c), consistent with findings that showed that solo-WCGW undergo replication-dependent methylation loss (63). In contrast, the PcG-marked CGI sites maintained similar methylation gain trajectories in cultures maintained with or without mitotic cycles (Fig. 7e). Thus, Polycomb-CGI methylation gains were not detectably coupled to the number of cell divisions under these experimental conditions and rather show a time-dependent increase. Together, these results support the model in which non-CGI, solo-WCGW hypomethylation is directly linked to replication history, whereas promoter-associated and Polycomb-marked CGI methylation gains reflect a distinct, time-dependent process independent of replication cycles.

Finally, we examined whether these patterns were also evident in vivo by analyzing methylation data from healthy colonic mucosa, tubular adenomas (76), and colorectal adenocarcinomas (COAD) from TCGA (Fig. 7g, h, Suppl. Fig. 5). Adenoma and COAD have lower methylation at the non-CGI regions compared to normal mucosa but showed similar methylation levels across ages (Fig. 7h). However, normal mucosa shows significant age-associated hypomethylation at solo-WCGW sites with increasing age (∼0.1% methylation loss per year, p < 0.01; Suppl. Fig. 5d), consistent with accumulation of replication-associated methylation loss during homeostatic turnover of the colonic epithelium. In adenomas and COAD, while solo-WCGW methylation was significantly lower than in healthy mucosa across the analyzed age range, the age-associated decline observed in healthy mucosa was absent. This pattern reflects methylation loss accumulated during multiple cell divisions during early stages of neoplastic expansion, followed by relative stabilization in the sampled lesions. In contrast, adenomas and COAD exhibited higher baseline methylation at PcG-associated CGIs than healthy mucosa across the age range. However, unlike solo-WCGW sites, PcG-associated CGIs continued to gain methylation with age, with broadly similar positive slopes across healthy mucosa, adenomas, and COAD (Fig. 7g). These data indicate an early increase in Polycomb-CGI methylation in neoplastic lesions that is superimposed on a disease-independent, shared age-associated trajectory of methylation gains. In summary, the in vivo data support observations made in the experimental models that CGI-methylation gains follow a time-dependent process distinct from the methylation loss at non-CGI regions represented by the solo-WCGW sites.

Collectively, these orthogonal analyses support the genomic context-dependent model of epigenetic remodeling, that is independent of oncogene-or mitogenic signaling. Polycomb-associated CGI methylation gains follow a shared, time-dependent trajectory across lineages and biological systems and are not detectably coupled to cell division or oncogenic signaling. This is in contrast to non-CGI sites that undergo replication-associated hypomethylation, accentuated in the *BRAF^V600E^* lineage. These findings suggest that promoter CGI hypermethylation, characteristic feature of cancers, is governed primarily by time-dependent epigenetic drift and chromatin context, distinct from the replication associated hypomethylation (62,63). This separation provides a framework wherein oncogenic-BRAF remodels the methylome, not by uniformly accelerating methylation change, but by selectively amplifying replication-sensitive hypomethylation against a background of shared CGI remodeling.

## Discussion

A long-standing question in cancer epigenetics is whether oncogenic signaling actively instructs de novo DNA methylation at CGI promoters, or whether oncogenes instead select for cells that have already acquired a permissive, age-associated methylation state. Because this distinction requires resolving methylation changes at clonal resolution and over physiologically relevant timescales, it has been difficult to test directly in vivo. We addressed this using an inducible *BRAF^V600E^* organoid model in which colonic stem cells undergo transformation over 5–6 months following oncogene activation, accompanied by methylation changes at key tumor suppressor genes that resemble those associated with aging (22). By combining a low-input, genome-wide eWGBS with longitudinal tracking of individual organoid clones, we show that *BRAF^V600E^* induction does not acutely increase the rate or magnitude of CGI methylation gain relative to isogenic, uninduced controls. Instead, both lineages (*BRAF^V600E^* and Control) accumulate methylation at the PcG-marked CGIs in a shared, time-dependent trajectory. In contrast, the *BRAF^V600E^* lineage selectively amplifies replication-associated methylation loss at non-CGI, late-replicating regions, which is distinctly observed at the solo-WCGW sites, consistent with the increased proliferation-associated hypomethylation at these sites (62,63). These results argue against a purely instructive model of BRAF-driven CGI methylation and instead support a model in which oncogenic-signaling acts upon, rather than generates, a pre-existing substrate of age-and drift-associated epigenetic change.

If *BRAF^V600E^* does not instruct new CGI methylation, what does it contribute? Our transcriptomic data suggest that its principal early effect is on cell-state selection rather than acute, de novo epigenetic reprogramming. scRNA-seq revealed a progressive, multi-month loss of goblet cell identity and accumulation of transit-amplifying and enterocyte-like populations in the *BRAF^V600E^* lineage, accompanied by downregulation of the intestinal stem and differentiation regulators such as *Cdx1*, *Cdx2*, *Gata4*, *Ascl2*, among the same regulators whose binding motifs are selectively enriched at *BRAF^V600E^*-specific differentially methylated sites. Although the direct role of these factors have to be established to define causative relationships, the data points to involvement of oncogene-induced transcriptional state differences in modulating PcG-CGI methylation, which progressively evolves over time. Mechanistically, PcG-CGI, which basally are lowly methylated and dependent on PRC2-associated repression rather than DNA methylation for their silencing, may be particularly susceptible to de novo methylation once repressive chromatin is established, regardless of the specific CpG site at which this occurs within the domain. *BRAF^V600E^*-driven alterations in lineage-restricting transcription factors could modulate the same stochastic, chromatin-context-governed methylation drift observed in control lineage to preferentially accumulate at a BRAF-relevant regulatory subset, without requiring a distinct, oncogene-specific methylation-writing mechanism. Supporting this selective rather than instructive role, *Cdkn2a* (*p16*) expression increased at one month, consistent with oncogene-induced senescence, then declined by five months. This decline paralleled the stepwise, progressive divergence of *BRAF^V600E^* clones in methylome space, indicating successive rounds of clonal selection/takeover by cells that bypass this senescence barrier. By contrast, control clones underwent stochastic drift followed by late convergence toward a stable epigenetic state, accompanied by upregulation of stress-and antiviral-response transcription factors (*Jun*, *Junb*, *Atf3*, *Irf9*, *Nfkb1/2*). The latter pattern is more consistent with adaptation to chronic culture stress than with oncogenic transformation. These parallel but distinct trajectories illustrate how a shared epigenetic substrate can give rise to divergent phenotypic outcomes depending on the selective pressures acting on it.

Our findings distinguish two genome domain-specific methylation trajectories with different relationships to proliferation. Proliferation-associated methylation loss at PMD-associated solo-WCGW sites is consistent with the established model of replication-coupled loss within partially methylated domains (62,63). In contrast, methylation gain at PcG-associated CGIs was decoupled from cumulative replication. We see this pattern across the organoid model, longitudinal cell line proliferation-perturbation experiments, and in human tissues across healthy colonic mucosa, tubular adenomas, and TCGA colorectal adenocarcinoma. Together, these observation across independent systems indicate that the decoupling of CGI hypermethylation from replication is not a peculiarity of our model but a generalizable feature of epigenetic remodeling during aging and tumorigenesis. To our knowledge, this is the first demonstration of this decoupling using longitudinal, clonal-resolution kinetic data rather than inference from cross-sectional comparisons, and it provides a mechanistic explanation for why CGI-hypermethylation and global hypomethylation, two key hallmarks but seemingly contradictory features of the cancer methylome, can co-occur within the same genome.

These observations suggest that PcG-CGI methylation gain captures time-dependent epigenetic remodeling rather than functioning exclusively as a counter of cell divisions. This distinction has implications for the mechanistic interpretation of mitotic clocks, such as epiTOC, which use methylation gain at selected Polycomb-associated promoter CpGs to infer proliferative history. Their development relied on mathematical modeling of age-associated methylation changes and comparisons with independently estimated tissue-specific stem-cell division rates, rather than direct longitudinal measurements of cell divisions and methylation changes at these sites. Our findings would challenge the specificity of these clocks for cumulative stem-cell divisions, without necessarily diminishing their predictive associations with aging or cancer risk (47,65,66). Possible explanation for the association between PcG-CGI methylation gain and tissue-specific division rates is that tissue-specific exposures to injury, inflammation, and regenerative signaling contribute to both proliferative activity and PcG-CGI methylation gain, producing an association without requiring methylation gain to be directly coupled to replication. Mechanistically, this possibility is supported by precedent that methylation of PcG target genes arise during inflammation (16,17,17,77,78), persistent DNA damage responses (20), environmental exposures (19) and metabolic modulation of epigenetic factors (79,80). Defining how these processes interact with chromatin context (e.g., Polycomb marking) and with clonal selection pressures imposed by specific oncogenic processes will be necessary to determine whether age-and drift-associated hypermethylation can be therapeutically slowed, prevented, or reversed, which is an important open question given that our data suggest this process begins well before, and independently of, oncogene-driven transformation.

There are several limitations to this study that need to be considered and addressed in future studies. Firstly, organoid models, while physiologically superior to cell-line models for capturing native epithelial heterogeneity and differentiation, still lacks stromal, immune, and microbial inputs that may potentially shape methylation dynamics in vivo. Thus, the degree to which culture-associated epigenetic drift quantitatively mimics the in vivo chronological aging remains an approximation, though its qualitative concordance with human mucosa, adenoma, and TCGA COAD data (Fig. 7g, h) supports our key conclusions. Second, the transcription factor– methylation associations are correlative. While the temporal coincidence of expression changes of several key TFs relevant to CRC biology, such as CDX1, CDX2, PPPARG, within *BRAF^V600E^*-specific methylation changes are consistent with their role in modulating the methylation drift, its causative roles needs to be functionally tested, i.e. whether restoring these factors prevents or reverses the associated methylation changes. Third, motif-enrichment and pseudobulk expression analyses cannot resolve whether transcription factor loss precedes, follows, or occurs independently of local methylation change at a given locus; single-cell multi-omic approaches capturing methylation and expression in the same cells would help resolve this causal ordering.

In summary, by resolving DNA methylation dynamics at clonal resolution over a physiologically relevant time course, our findings reframe the relationship between oncogenic signaling and promoter CpG-island hypermethylation. Rather than instructing de novo methylation gain, *BRAF^V600E^* acts within a shared, time-dependent substrate of CGI methylation drift. Replication-associated methylation loss at the solo-WCGW sites within partially methylated domains robustly reflects cell division history, as demonstrated by a previous study (63). This decoupling of CGI hypermethylation from both replication and oncogenic signaling identifies chromatin context and time-dependent drift as the primary driver of this hallmark cancer epigenetic lesion. More broadly, this work provides a conceptual framework and methodological strategies for dissecting how clonal selection operates on pre-existing epigenetic heterogeneity during the earliest stages of tumor evolution. These findings have implications both for how promoter methylation is interpreted as a biomarker, and for identifying the upstream physiological processes that generate this epigenetic substrate as potential targets for early intervention.

## Methods

### Organoid culture

Male mice were euthanized via Carbon Dioxide, the first 2.5 cm of the proximal colon was collected, colon crypts were isolated and the organoid culture was established as described previously (81,82). Organoids were plated within Matrigel (Corning #356231). Culture media was comprised of 50% of Wnt3a conditioned media, 20% of R-Spondin 1 conditioned media, 20% of Advanced DMEM/F12 (ThermoFisher #12634010) with B27® (ThermoFisher #17504044), 10% of Noggin conditioned media and EGF. The conditioned media for Wnt3a, R-Spondin1, and Noggin were made as described previously (83). Lentiviral delivery of Cre was used subsequently to activate Braf^V600E^ within the organoids, infected organoids were selected with a Braf^V600E^ mutation with either 200 µg/ml hygromycin or full medium without EGF for two weeks. The Braf^V600E^ and EV induced organoids were grown for five months. Each month the organoids were dissociated to single cell and replated. The single cells were grown for 8-10 days to form single organoids from single cells. Each month 5 single organoids were picked for eWGBS.

### Dissociation of Organoid to single cell

The organoids were dissociated to single cell via Dispase and Accumax. Briefly, the organoids were rinsed using PBS and 2 ml of Dispase was added. The organoids were disrupted by using a cell scraper and broken up by pipetting up and down for 10-15 times. The organoids were incubated at 37°C for 15 mins and centrifuged at 398 g for 5 mins. The organoids were resuspended with PBS by pipetting up and down for 10-15 times and centrifuged at 398 g for 5 mins. 2 ml of Accumax was added and mixed, further incubated for another 15 mins at 37°C. The organoids were centrifuged at 398 g and resuspended with 1 ml of PBS by pipetting up and down 10-15 times. Further, the cells were dispersed in a 21-G needle and passed to a 20 μM (CellTrics) filter to obtain the single-cell population.

### Subcloning of Lentivirus Plasmid and Packaging of Lentivirus

Subcloning of Lentivirus Plasmid was done as described previously (22). Lentivirus was produced in Lenti-X™ 293T cell line transfected with Packaging vectors and Lenti-cre or Lenti-empty mixed with Xfect™ polymer. After 72 hrs of the transfection, the media was collected and concentrated using Lenti-X™ concentrator.

### DNA Isolation

DNA isolation was done by using the Quick-DNA Microprep Kit (ZYMO RESEARCH #D3020) as per the manufacturer’s instructions with few modifications. Briefly, single organoids were lysed by using the 50 µl lysis buffer. Further, the single organoids were resuspended by vertexing for 6 seconds and incubated for 5-10 minutes at room temperature. The mixture was transferred to a Zymo-Spin column and centrifuged at ≥ 10,000 x g for one minute. The flow through was discarded and 500 µl of g-DNA wash buffer was added to the spin column and centrifuged at ≥ 10,000 x g for one minute. Further, 200 µl of DNA pre-wash buffer was added to the spin column and centrifuged at ≥ 10,000 x g for one minute. The spin column was transferred to a microcentrifuge tube and 12 µl DNA elution Buffer was added followed by centrifugation at ≥ 12,000 x g for 30 seconds to elute the DNA.

### Bisulfite Conversion

Bisulfite conversion was performed on DNA from single organoid by using EZ DNA Methylation-Direct kit (ZYMO RESEARCH #D5020) according to the manufacturer’s protocol with the following modification. 65 µl of CT conversion reagent was added to the 10 µl of DNA.15 *fg* of lambda DNA was also added as a spike-in control. The bisulfite conversion was performed by using the following cycle in a thermocycler 98°C for 8 minutes, 64°C for 3.5 hours,4°C for hold. Further, 300 µl of M-Binding buffer and 3 µg Carrier RNA was added to the bisulfite-converted DNA and mixed it well. The mixture was transferred to a Zymo-Spin column and centrifuged at ≥ 12,000 x g for one minute and the flow through was discarded. The columns were further washed by using 100 µl M-Wash buffer and centrifuged at (16,000 x g) for 1 minute. For desulphonation, 200 µl of M-Desulphonation buffer was added to the column and let stand for 20 mins at room temperature. The tubes were centrifuged at (16,000 x g) for 1 minute. The columns were washed 2 times by using M-Wash buffer and for the elution 12 µl of elution buffer was added to the column to purify the DNA.

### Emulsion Multiple Displacement amplification

MDA was performed using bisulfite-converted DNA. The MDA reactions were set up by using 1 X Phi29 buffer, 2.8 mM DNTPs, 1.1% tween 40, 24 mM of bisulfite-aware DNA primers (7 *mer* without C and G), 12 mM of random 7 *mer*, 2 ng/μl sonicated poly-adenylic acid (poly-A), 15 mM trehalose dihydrate, 2mg BSA and H2O in a single tube. The reaction was incubated at 95°C for 1 min followed by cooling to 4°C and transferred to Ice for 2 mins. 1µl (100U/μl) Phi 29 DNA polymerase and 1µl (50 U/ul) of Klenow fragment exo-(Enzo) were added and mixed the 50 µl of reaction into the 300 µl oil in 1.5 ml low DNA binding tube and further vortex it for 4 mins in the cold room (4°). Aliquoted 100 µl each in PCR tubes and incubate the reactions at +4°C/15s to 37°C, 37°C for 17 hrs followed by a 10-min incubation at 85°C. For breaking the emulsion and purification Micellula DNA emulsion & purification kit was used (EURx #3600). The purification was done by using the manufacturer’s instructions.

### Library Preparation

Library preparation was done by using the KAPA Hyper Prep Kit as per the manufacturer’s protocol with some modifications. Briefly, end repair and A-tailing were performed by taking the fragmented (200-300 bp) equal amount of double standard DNA followed by adapter ligation by using the (IDT-TrueSeq DNA UD) dual index (Illumina #20022370). Post-ligation clean-up was done by using AMPure XP beads (Beckman, # A63881) with 0.8x bead solution to DNA volume was performed. Purified DNA fragments were taken for library amplification. The adapter-specific primers were used for library amplification. The amplified libraries were purified by using the 1X AMPure XP beads solution. Further libraries were quantified by using the Kapa Library Quantification kit (Biorad #KK4844). To determine the size distribution Agilent Bioanalyzer was used by using the Agilent High Sensitivity DNA kit (Agilent, #50674626). Further, the libraries were pooled to a concentration of 4 nM and sequenced on the Illumina NovaSeq 6000. Demultiplexing and FASTQ generation were completed using Illumina’s BaseSpace software.

### Single cell RNA sequencing library prep

The BrafV600E and EV-induced organoids were grown for five months and each month the organoids were dissociated to single cell by using the enzymatic method (Dispase+Accumax). Cell counts and viability were determined using the Cell Countess II with Trypan Blue. A maximum volume of 43.3 uL/sample was used for processing to target up to 10,000 cells. Cells were combined with RT reagents and loaded onto 10X Next GEM Chip G along with 3’ v3.1 gel beads. The NextGEM protocol was run on the 10X Chromium Controller to create GEMs, composed of a single cell, gel bead with unique barcode and UMI primer, and RT reagents. 100uL of emulsion is retrieved from the chip and incubated (45 min at 53C, 5 min at 85C, cool to 4C), generating barcoded cDNA from each cell. The GEMs are broken using Recovery Agent and cDNA is cleaned, following the manufacturer’s instructions using MyOne SILANE beads. cDNA is amplified for 11 cycles (3 min @ 98C, 11 cycles: 15sec @98C, 20sec @63C, 1min @72C; 1min @ 72C, cool to 4C). Samples are cleaned using 0.6X SPRI select beads. QC was completed using Qubit and Bioanalyzer to determine DNA size and concentration. 10uL of amplified cDNA was carried into library prep. Fragmentation, end repair and A-tailing were completed (5 min @ 32C, 30 min @ 65C, cool to 4C), and samples were cleaned using double-sided size selection (0.6X, 0.8X) with SPRI select beads. Adaptor ligation (15min @ 20C, cool to 4C), 0.8X cleanup, and amplification were performed via PCR using unique i7 index sequences. Libraries underwent a final cleanup using double sided size selection (0.6X, 0.8X) with SPRI select beads. Library QC was performed via Qubit, Bioanalyzer, and KAPA library quantification qPCR kit. Libraries were sequenced on the Illumina NovaSeq 6000 using v1.5 kits, targeting 50K reads/cell, at read lengths of 28 (R1), 8 (i7), 91 (R2). Demultiplexing and FASTQ generation were completed using Illumina’s BaseSpace software.

### Bioinformatic methods

eWGBS raw data were processed with FastQC (v0.12.1), adapters were trimmed using TrimGalore (v0.6.10) to remove the first 9 base pairs, and poor-quality reads, reads were mapped using Biscuit (v1.0.0.20210917) to mm10. Differential analysis was performed using limma via the RnBeads (v2.27.2) package. A CpG site was considered differentially methylated if it had an absolute delta methylation between the control and experimental states of greater than 0.2 (on a methylation scale of between 0 and 1; one being fully methylated, 0 being fully unmethylated) and an FDR adjusted p-value of less than 0.05.

### Analysis of The Cancer Genome Atlas data

Level 1 The Cancer Genome Atlas (TCGA) Illumina HumanMethylation450 BeadChip Array data were downloaded from TCGA for colon adenocarcinoma (COAD). Data were loaded into R 4.5.1 (RRID: SCR_001905), preprocessed to remove all probes on the X and Y chromosomes due to potential cross hybridization risks. Probes which reside on SNPs were removed, and those with detection P values > 0.05 were removed. Probes on the Illumina probe blacklist for the HumanMethylation450 BeadChip Array were also removed (84). Data were normalized using functional normalization and (Minfi v1.55.1, RRID: SCR_012830) combined with tubular adenoma data from (76).

### KEGG analysis

Briefly, differentially methylated sites which satisfied the criteria mentioned in the above paragraph were split into hyper/hypomethylated sites as well as into their associated genomic element (promoters with CGI, promoters without CGI). A promoter was defined as a TSS +/- 1500bp and CGI definitions were taken from UCSC genome browser mm10. Gene symbols associated with the CG sites were then converted into EntrezIDs via AnnotationDbi (v1.72.0) and KEGG enrichment analysis was computed using the enrichKEGG() from clusterProfiler (v4.20.1) with a p value cutoff of 0.05 and the Benjamini Hochberg (BH) method for p value adjustment. Results were plotted via dotplot().

### Transcription factor enrichment analysis

Differentially methylated sites from RnBeads (v2.27.2) were obtained for *BRAF^V600E^*5-month vs 1-month and control 5-month vs 1-month and subset to promoters with CGI. A promoter was defined as a TSS +/- 1500bp and CGI definitions were taken from UCSC genome browser GRCm38/mm10. Given the output from differential methylation would be one CG site in length and to enable transcription factor motif binding enrichment within the area, differential sites were given 50bp flanks. Any resulting region which overlapped with another either within or between lineages was excluded from any downstream transcription factor enrichment analysis. Transcription factor enrichment analysis was facilitated via SEA (Simple Enrichment analysis) via the MEME Suite (v5.5.9) using the HOCOMOCO H12 Core database. Statistically significant output transcription factors were then compared between lineages to obtain transcription factor profiles unique to each lineage.

### Single cell RNA-seq

scRNA-seq raw data were processed using Cell Ranger 6.0.1 (10X Genomics) with default parameters aligned to mm10 and then analyzed using Seurat (v5.3.0). Each sample was read in to create a separate layer of a Seurat object, labelled additionally by treatment vector (*BRAF^V600E^* or Control). Genes that did not occur in a minimum of 10 cells were removed and cells that did not have a minimum of 200 features were removed. Mitochondrial reads were also regressed out within SCTransform(). Layers of the Seurat object were integrated via Harmony. Post integrative analysis with Harmony, layers were rejoined, neighbors found using the first 10 dimensions, clusters found using resolution 1 and markers found using FindAllMarkers().

Cell types were annotated via markers output from FindAllMarkers() within Seurat. Briefly, gene symbols output were converted to EntrezIDs using AnnotationDbi (v1.72.0) and org.Mm.eg.db (v3.22.0). Following this, EntrezIDs and their average Log2 fold change were grouped by cluster,sorted in descending order and input into easyct() from EasyCellType (v1.13.0) using the CellMarkers database, the GSEA method and the tissue type identifiers “Colon”, “Colon epithelium”, “Colorectum”, “Intestinal crypt” and “Intestine”. Identifiers were checked for congruency using cell marker 2.0 (85) missing identifiers were resolved using this database also.

Pseudobulk analysis was performed using DESeq2 (v1.4.3). Briefly, summed counts for each sample were obtained via AggregateExpression() then a DESeq object was created from the resulting matrix followed by DESeq() for normalization and differential expression analysis. Heatmaps were plotted using z scores resulting from a scaled variance stabilizing transformation on the normalized DESeq() output.

### Genomic Annotation

A promoter was defined as a transcription start site +/- 1500bp, CGI definitions were taken from UCSC genome browser for mouse GRCm38/mm10. Genomic intervals of polycomb marked CGI were obtained from Easwaran and colleagues (9).

## Data Availability

eWGBS and scRNA-seq data resulting from this investigation have been deposited in the Sequence Read Archive accession number PRJNA1552217.

## Acknowledgements

National Cancer Institute under award numbers R01CA229240, R01CA230995 (to H.E., and S.B.B.,), R21CA212495 (to H,E., T.P.); grant from the National Institute of Environmental Health Sciences, National Institutes of Health Institute under award number R01ES011858 (to S.B.B., H.E.,); the Evelyn Grollman Glick Scholar Award (H.E.,); and the National Cancer Institute under award number P30CA006973 (SKCCC Core Grant); funding from the Schnabl Foundation. We thank members of our laboratories for critical review of the manuscript, Tina Largent and Ray-Whay Chiu Yen for lab support and JoAnn Johnson, Lauren Bois, and Tammy Means for administrative support. The content is solely the responsibility of the authors and does not necessarily represent the official views of the National Institutes of Health or other funders.

## Supplemental Legends

**Supplemental Figure 1:**
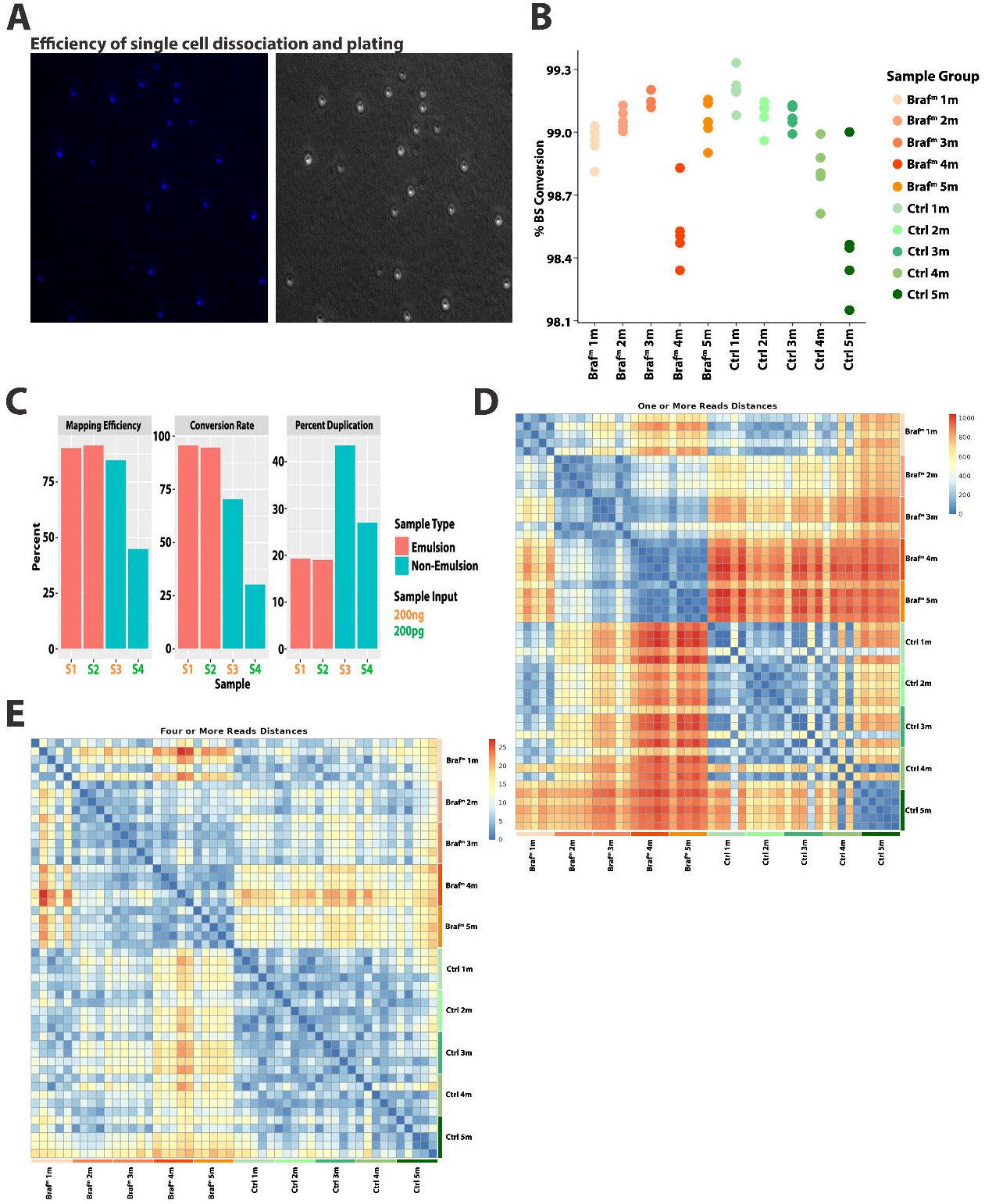
Technical Validation and Quality Assessment of Low-Input Emulsion-Based WGBS Profiling of Single-Cell-Derived Organoid Clones. (A) Single-cell dissociation following treatment with Dispase and Accumax. Representative fluorescence (left) and transmitted-light (right) images of the resulting cell suspension show predominantly individual cells, with DAPI visible in the left panel. (B) Percent bisulfite conversion rate across control (ctrl) and *BRAF^V600E^* (Braf^m^) lineages colored by lineage and timepoint. (C) Percent mapping efficiency, conversion rate and percent duplication of emulsion (red) and non-emulsion (blue) based amplification. Sample names colored by sample input amount, 200pg (orange), 200pg (green). (D-E) Euclidean distance matrices between all clones within regions with one or more reads (D) and four or more reads (E).

**Supplemental Figure 2:**
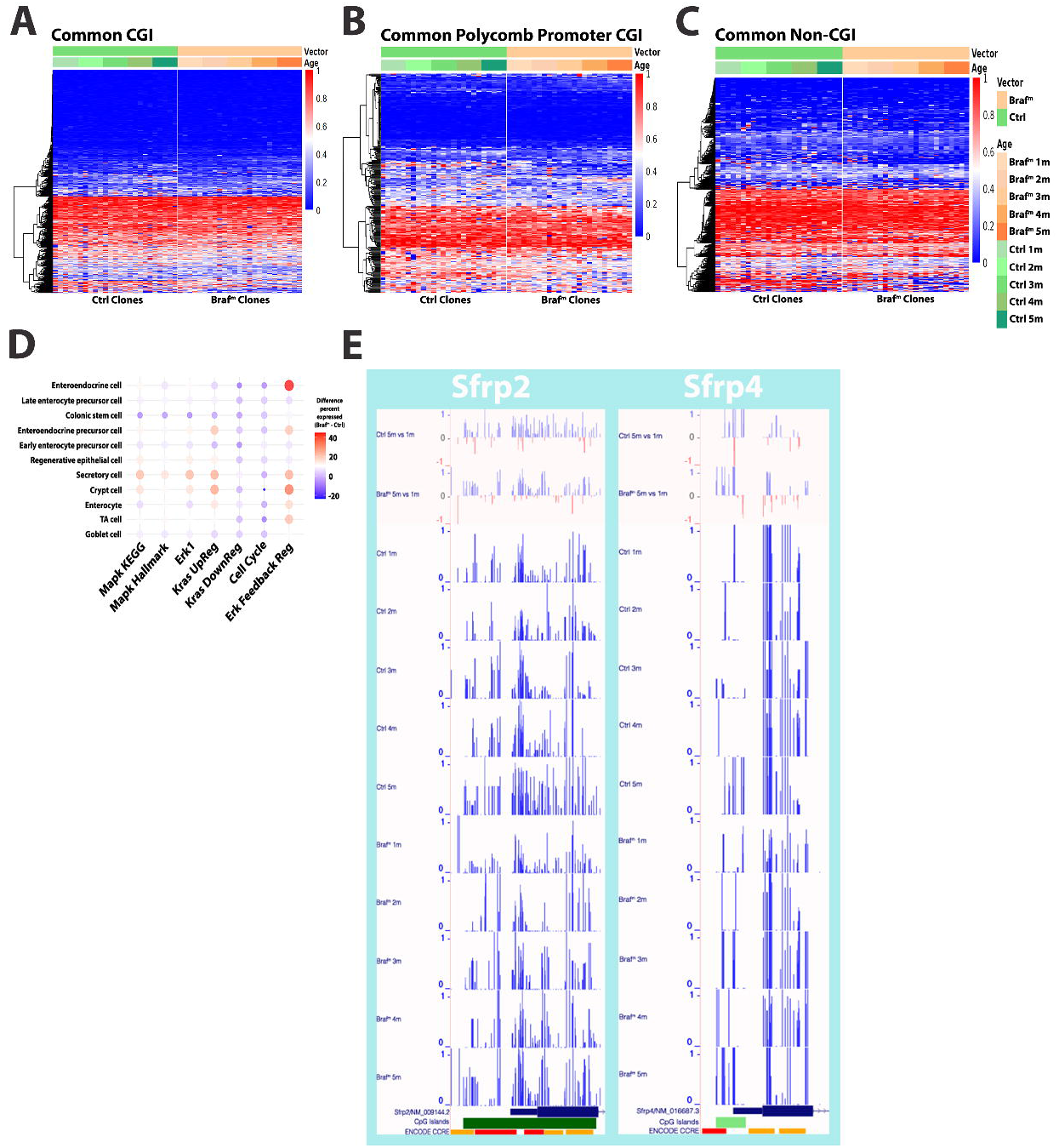
Longitudinal DNA Methylation Profiles and MAPK-Associated Transcriptional Signatures in BRAFV600E mutant and Control Organoid Clones. (A-C) Heatmap with unsupervised hierarchical row clustering colored by control (Ctrl) and *BRAF^V600E^* (Braf^m^) across different genomic elements CGI (A), polycomb marked promoter CGI (B), non-CGI sites (C). (D) Difference in percent expression (scRNA-seq) across different genes, pathways and cell types between control (Ctrl) and *BRAF^V600E^*(Braf^m^) lineages. (E) UCSC Genome Browser tracks showing the difference in methylation (ΔMeth-orange shaded boxes) and absolute methylation for each timepoint and lineage across genes which have been observed to show promoter hypermethylation during colorectal cancer.

**Supplemental Figure 3:**
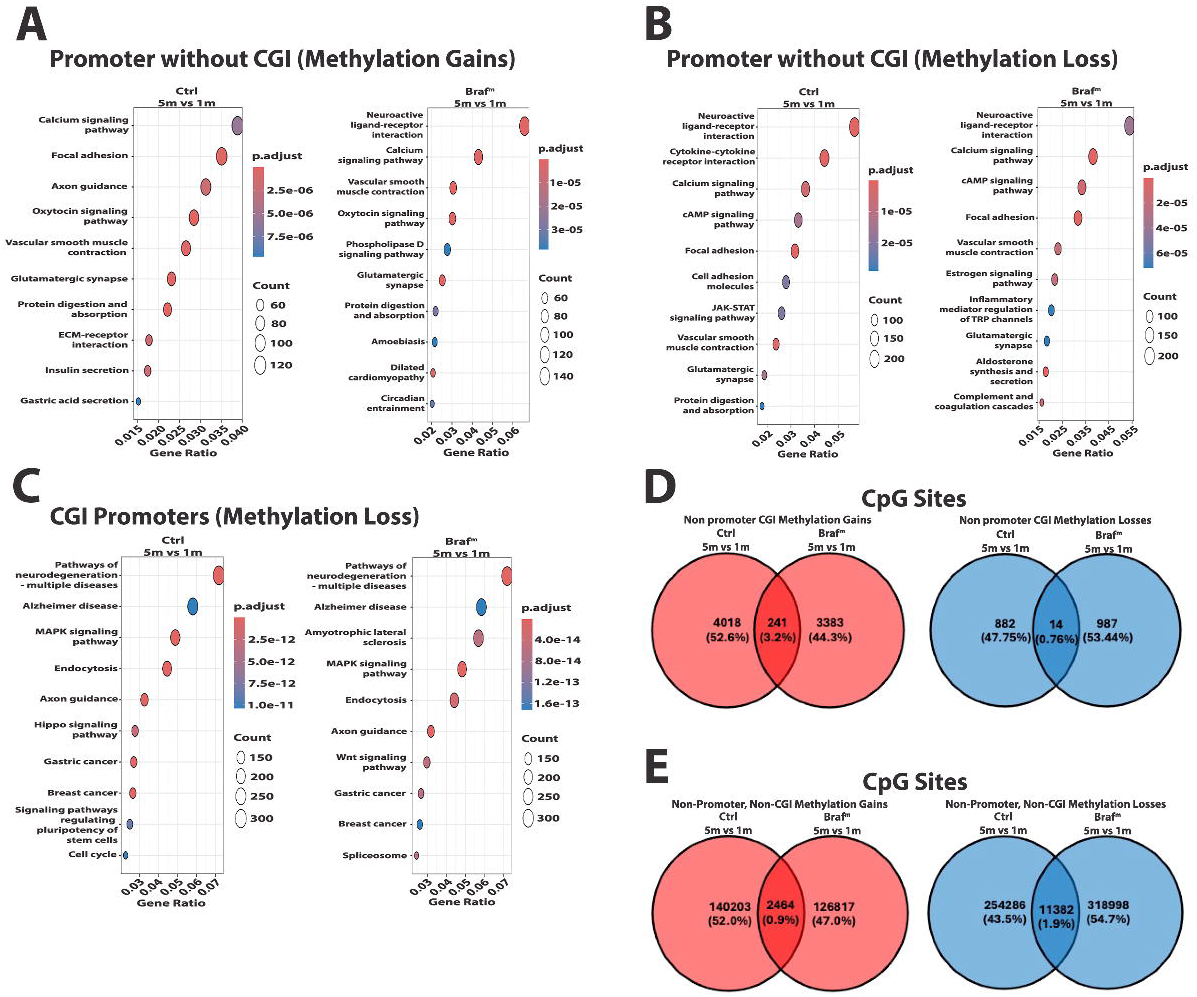
Genomic distribution, pathway enrichment, and transcription factor–associated features of longitudinal DNA methylation changes in control and *BRAF^V600E^* organoids. DNA methylation changes were assessed between months 1 and 5 within control (Ctrl) and *BRAF^V600E^*(Braf) organoid lineages. (A, B) Pathway enrichment analyses of genes associated with promoters lacking CpG islands (CGIs) that gain methylation (A) or lose methylation (B). Control and *BRAF^V600E^* results are shown on the left and right, respectively. (C) Pathway enrichment analyses of genes associated with CGI-containing promoters that lose methylation in control (left) and *BRAF^V600E^*(right) organoids. (D, E) Venn diagrams showing shared and lineage-specific methylation gains (red) and losses (blue) at non-promoter CGI sites (D) and CpG sites outside both promoters and CGIs (E).

**Supplemental Figure 4:**
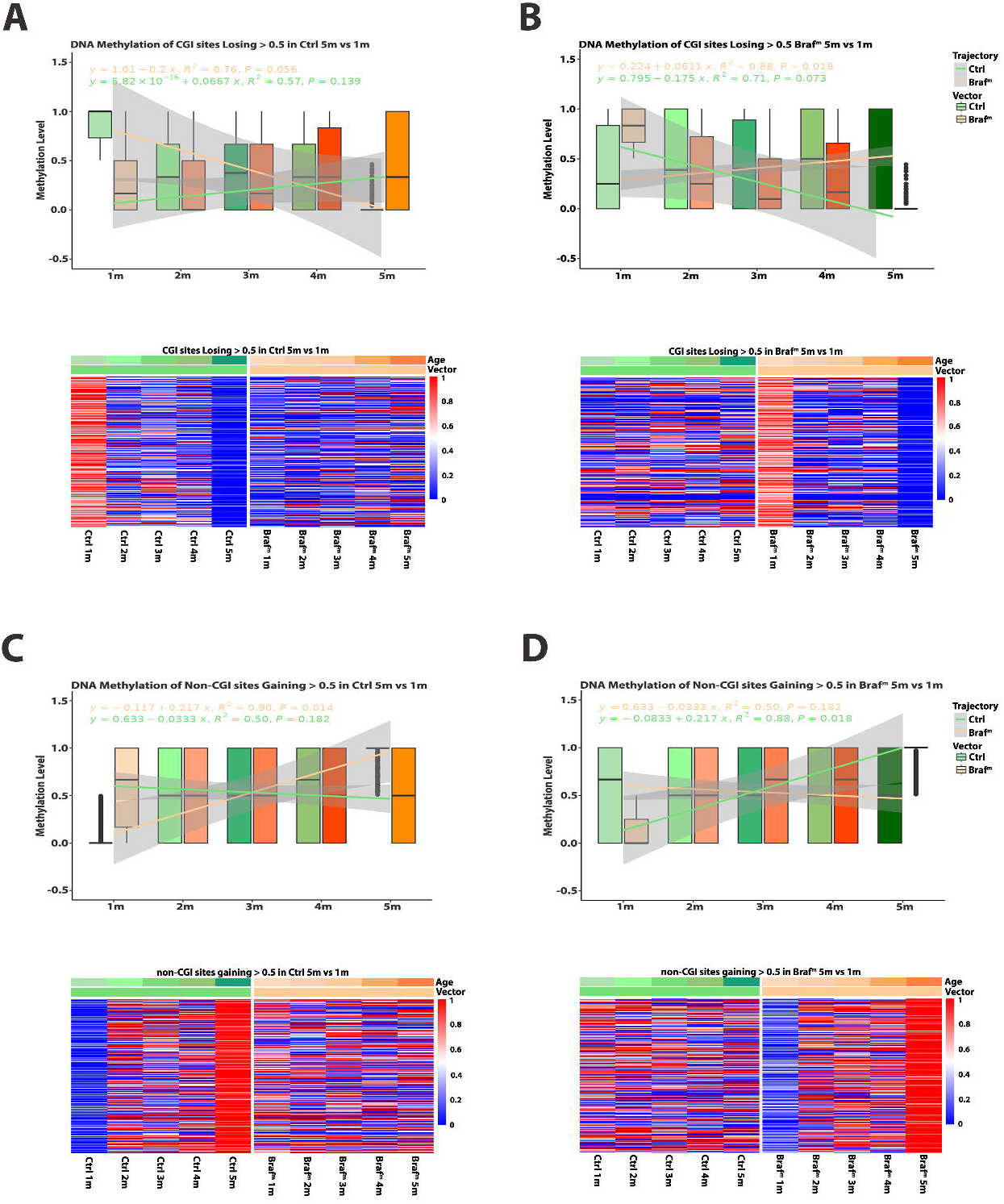
Longitudinal analysis of the DNA methylation at different genomic elements. (A, B) CGI sites losing (ΔMeth ≤ 0.5) in control (Ctrl) (A) or *BRAF^V600E^*(Braf^m^) (B) across lineages and month timepoints. (C, D) Non-CGI sites gaining (ΔMeth ≥ 0.5) in control (C) and *BRAF^V600E^*(D) across lineage and month timepoints. Heatmap below each boxplot shows the methylation of all selected genomic element related CpG sites across lineages and time. Selection criteria of the sites investigated can be observed in the plot title. Boxplots colored by lineage (orange – *BRAF^V600E^*, green – Control) and timepoint, lines represent a smoothed linear model with equations, R^2^ and p values noted per lineage.

**Supplemental Figure 5:**
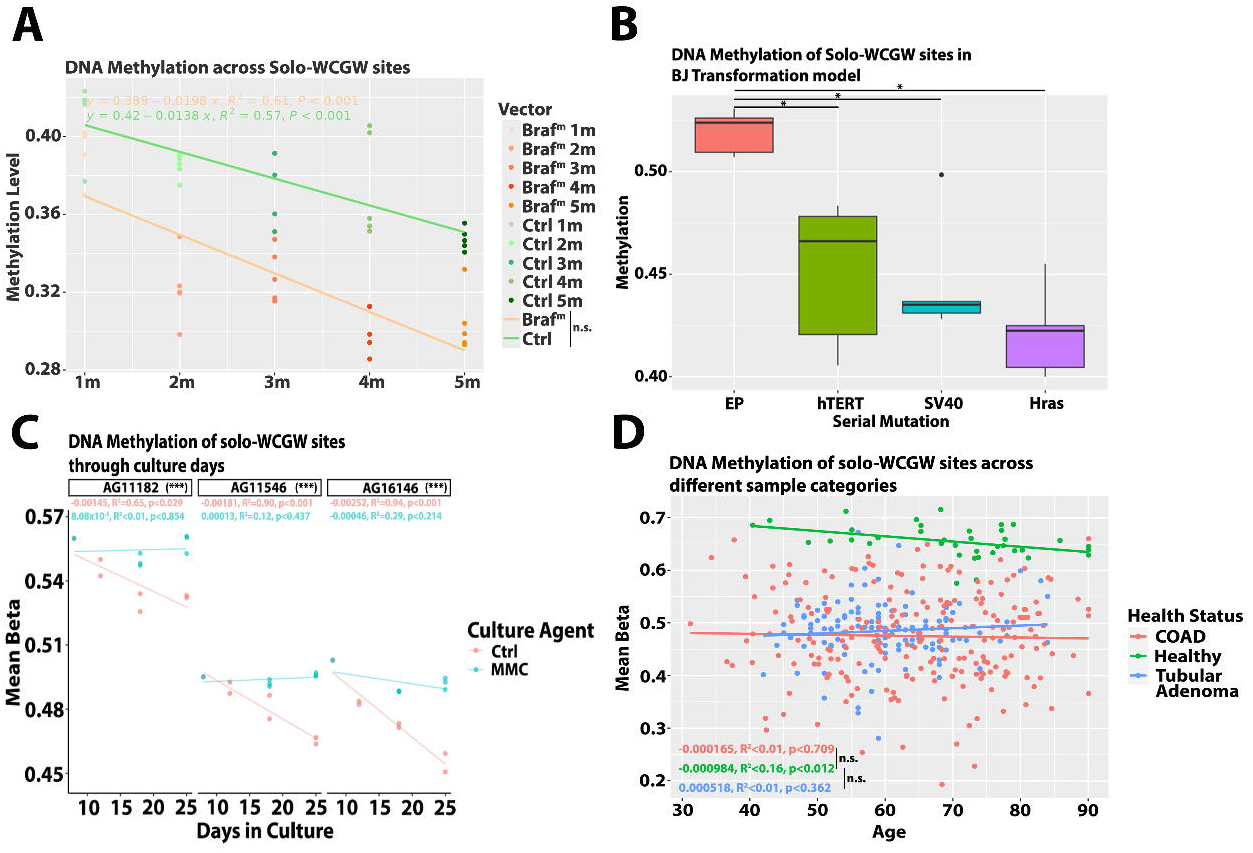
Replication-Dependent DNA Methylation Loss at Solo-WCGW Sites Is Retained Across Experimental Models. (A) Longitudinal analysis of the DNA methylation of Solo-WCGW sites in control (Ctrl, green) and *BRAF^V600E^* (Braf^m^, orange) lineages across 1-month to 5-month timepoints. Individual organoid clones colored by lineage (orange – *BRAF^V600E^*, green – Control) and timepoint, lines represent a smoothed linear model with equations, R^2^ and p values noted per lineage. (B) DNA methylation of Solo-WCGW sites upon sequential addition of proliferative agents, data from GSE91069. (C) Investigation of DNA methylation of Solo WCGW sites across culture days within different cell lines treated with Mitomycin C (MMC) or control treatments. Cell line identifiers noted in the graph facet title, interaction p-value noted in brackets within the facet title, linear model slope values, R^2^ and associated linear model p-value colored per intervention. (D) Investigation of DNA methylation in Solo-WCGW sites in TGCA COAD and tubular adenoma from GSE233616. Lines represent linear models with slope values, R^2^ and p-values colored per health state. P-values relating to interactions between models are shown via a vertical line listed in the graph legend. N.s. (p>0.05), * (≤0.05), ** (≤0.01), *** (≤0.001), **** (≤0.0001). Wilcox signed rank test unless otherwise specified. Abbreviations: EP – Early Passage, COAD – Colon adenocarcinoma.

## Notes

### Competing Interest Statement

The authors have declared no competing interest.

